# Structural role of essential light chains in the apicomplexan glideosome

**DOI:** 10.1101/867499

**Authors:** Samuel Pazicky, Karthikeyan Dhamotharan, Karol Kaszuba, Haydyn Mertens, Tim Gilberger, Dmitri Svergun, Jan Kosinski, Ulrich Weininger, Christian Löw

**Affiliations:** Centre for Structural Systems Biology (CSSB), Notkestrasse 85, D-22607 Hamburg, Germany; Molecular Biology Laboratory (EMBL), Hamburg Unit c/o Deutsches Elektronen Synchrotron (DESY), Notkestrasse 85, D-22607 Hamburg, Germany; Bernhard Nocht Institute for Tropical Medicine, Bernhard-Nocht-Strasse 74, D-20359 Hamburg, Germany; Department of Biology, University of Hamburg, Hamburg, Germany; Structural and Computational Biology Unit, European Molecular Biology Laboratory, Meyerhofstrasse 1, 69117 Heidelberg, Germany; Martin-Luther-University Halle-Wittenberg, Institute of Physics, Biophysics, D-06120 Halle (Saale), Germany

## Abstract

Apicomplexan parasites, such as *Plasmodium falciparum* and *Toxoplasma gondii*, traverse the host tissues and invade the host cells exhibiting a specific type of motility called gliding. The molecular mechanism of gliding lies in the actin-myosin motor localized to the intermembrane space between the plasma membrane and inner membrane complex (IMC) of the parasites. Myosin A (MyoA) is a part of the glideosome, a large multi-protein complex, which is anchored in the outer membrane of the IMC. MyoA is bound to the proximal essential light chain (ELC) and distal myosin light chain (MLC1), which further interact with the glideosome associated proteins GAP40, GAP45 and GAP50. Whereas structures of several individual glideosome components and small dimeric complexes have been solved, structural information concerning the interaction of larger glideosome subunits and their role in glideosome function still remains to be elucidated. Here, we present structures of a *T. gondii* trimeric glideosome sub complex composed of a myosin A light chain domain with bound MLC1 and TgELC1 or TgELC2. Regardless of the differences between the secondary structure content observed for free *P. falciparum* PfELC and *T. gondii* TgELC1 or TgELC2, the proteins interact with a conserved region of TgMyoA to form structurally conserved complexes. Upon interaction, the essential light chains undergo contraction and induce α-helical structure in the myosin A C-terminus, stiffening the myosin lever arm. The complex formation is further stabilized through binding of a single calcium ion to *T. gondii* ELCs. Our work provides an important step towards the structural understanding of the entire glideosome and uncovering the role of its members in parasite motility and invasion.

**Author summary:** Apicomplexans, such as *Toxoplasma gondii* or the malaria agent *Plasmodium falciparum,* are small unicellular parasites that cause serious diseases in humans and other animals. These parasites move and infect the host cells by a unique type of motility called gliding. Gliding is empowered by an actin-myosin molecular motor located at the periphery of the parasites. Myosin interacts with additional proteins such as essential light chains to form the glideosome, a large protein assembly that anchors myosin in the inner membrane complex. Unfortunately, our understanding of the glideosome is insufficient because we lack the necessary structural information. Here we describe the first structures of trimeric glideosome sub complexes of *T. gondii* myosin A bound to two different light chain combinations, which show that *T. gondii* and *P. falciparum* form structurally conserved complexes. With an additional calcium-free complex structure, we demonstrate that calcium binding does not change the formation of the complexes, although it provides them with substantial stability. With additional data, we propose that the role of the essential light chains is to enhance myosin performance by inducing secondary structure in the C-terminus of myosin A. Our work represents an important step in unveiling the gliding mechanism of apicomplexan parasites.

## Introduction

*Apicomplexa* are a phylum of intracellular, parasitic, single cell eukaryotes with a high medical and agricultural relevance. For instance, *Plasmodium species* is the causative agent of malaria, that leads to 414.000 deaths per year [1]. The number of malaria cases increased or stopped decreasing in several African countries in the last few years due to the emergence of new drug-resistant strains [1]. Another apicomplexan parasite, *Toxoplasma gondii*, is responsible for toxoplasmosis in humans [2]. Although more than 30% of the world population is thought to be infected with *T. gondii* causing no obvious symptoms, the infections can cause severe damage in immunocompromised patients and pregnant women [2]. Proliferation and transmission of these obligate endoparasites in theirs host organisms rely on efficient cell invasion [3]. This active process is based on the motility of the parasite that is referred as gliding and is empowered by an actin/myosin motor [4, 5]. This motor is localized within the intermembrane space between the parasite’s plasma membrane and inner membrane complex (IMC), an additional double-layer of membranes that is unique for these single cell organisms [6]. The IMC provides stability to invasion competent stages of the parasite and functions as an anchor for the actin/myosin motor. While motility is achieved by the interaction of the myosin with actin filaments, the myosin is linked to the IMC by a membrane-embedded multi-protein complex referred to as the glideosome [7–9] (Fig 1A).

**Fig 1.**
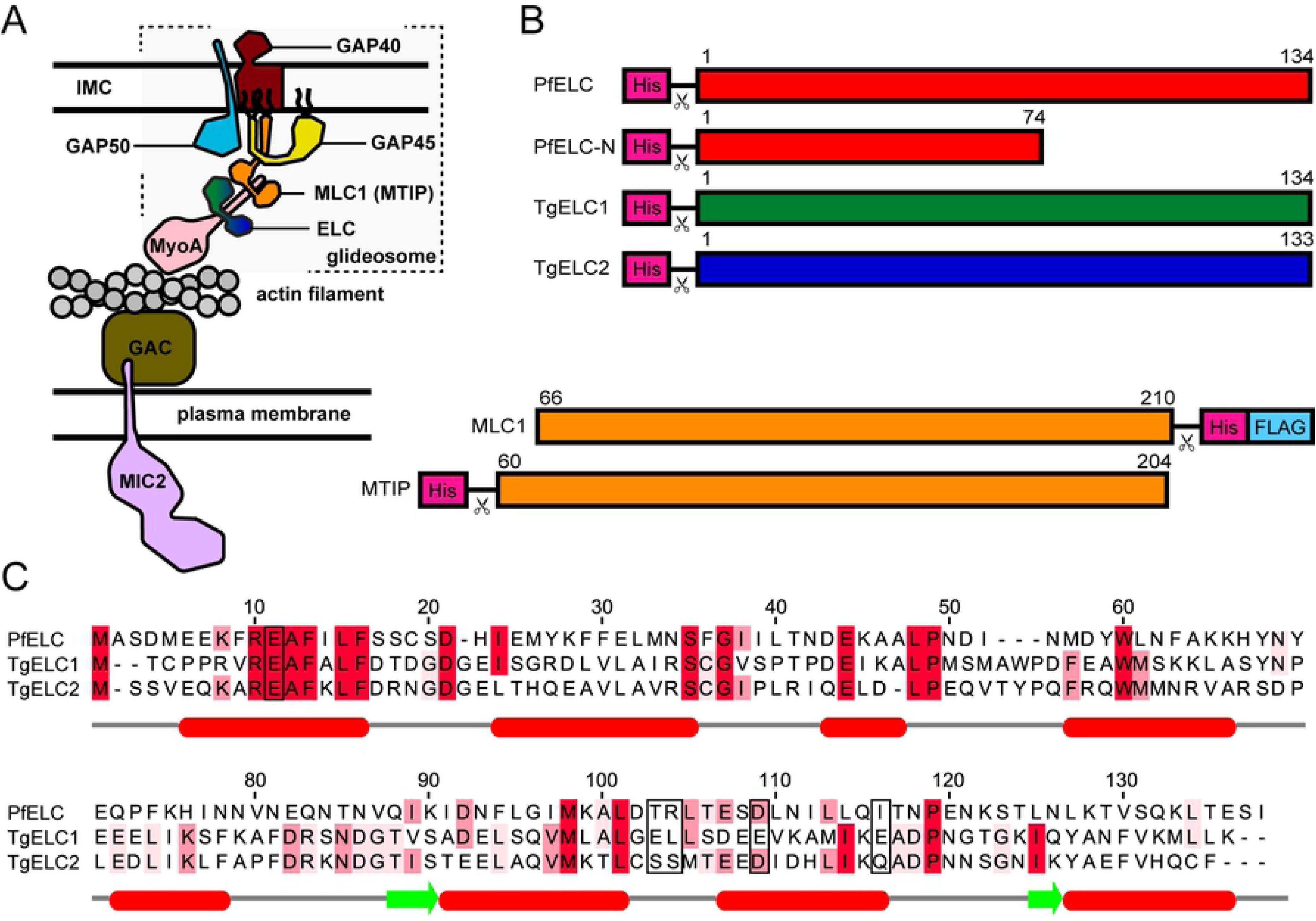
Glideosome and myosin light chains. (A) Schematic representation of the current model of the glideosome and its localization in the parasites intermembrane space. Actin is immobilized to the plasma membrane whereas myosin A is part of the glideosome, which binds the essential light chains ELC and myosin light chain MLC1 (called myosin tail interacting protein, MTIP, in *Plasmodium spp.*). Myosin A and its light chains further interact with glideosome associated proteins GAP40, GAP45 and GAP50, which anchor the glideosome in the outer membrane of the inner membrane complex. (B) Schematic representation of the myosin light chain constructs used in this study. The numbers indicate the sequence residues of the particular protein; the scissor symbol represents a TEV cleavage site. (C) Sequence alignment of *P. falciparum* PfELC and *T. gondii* TgELC1 and TgELC2. Identical residues between these proteins are highlighted in red. The boxed residues indicate the residues involved in the polar interactions with TgMyoA (see Fig 5 and S4 Table). Secondary structure elements of PfELC as predicted by JPred are graphically shown under the sequence alignment.

According to the current model, the apicomplexan glideosome is composed of six proteins: myosin MyoA, essential light chain ELC, myosin light chain MLC1, and the glideosome-associated proteins GAP40, GAP45 and GAP50 [7,8,10]. MyoA is an unusual small myosin protein of the unconventional myosin class XIV [11, 12], missing the typical myosin tail domain and binding the two light chains at the C-terminal myosin neck region [13, 14]. MLC1 (in *P. falciparum*: myosin A tail-interacting protein, MTIP) binds at the very C-terminus of MyoA, while ELC is expected to interact with the C-terminus of MyoA upstream of MLC1 [15]. Two ELC homologs recognizing the same MyoA region, termed TgELC1 and TgELC2, were identified in *T. gondii* [16], whereas only one PfELC homolog is known in *P. falciparum* [14, 17]. Myosin together with its light chains and the glideosome associated protein 45 (GAP45) has been shown to form a pre-complex in the earlier stages of intracellular parasite development [7], which subsequently assembles with the remaining glideosome members (GAP40 and GAP50). Both MLC1 (MTIP) and GAP45 use their N-terminal myristoylation and palmitoylation sites to anchor in the outer IMC membrane [18]. GAP45 is essential for the correct localization of MyoA with its light chains and GAP45 depletion leads to impairment of the host cell invasion [10]. Depletion of GAP40 or GAP50 changes the morphology of the parasites and the integrity of the IMC and thereby also alters the localization of MyoA and the light chains [19]. Thus, GAPs do not only serve as an anchor of the glideosome but provide stability and integrity to the IMC. Indeed, the localization of GAP40 is restrained to distinct foci evenly distributed along the co-localized tubulin, suggesting that the glideosome is further interconnected within the network of IMC proteins and in turn attaches to the cytosolic tubulin network [19, 20].

Structural information on individual members and subcomplexes of the glideosome are limited and the architecture of the entire glideosome is elusive. So far, only the structures of *P. falciparum* PfGAP50 soluble domain [21], a *T. gondii* dimeric complex between the TgMyoA C-terminus and MLC1 [15], a homologous dimeric complex in *P. falciparum* between PfMyoA C-terminus and MTIP [22], and the motor domains of the *T. gondii* TgMyoA [23] and *P. falciparum* PfMyoA [24] are available (S1 Table).

Here we present crystal structures of trimeric complexes of *T. gondii* composed of MLC1, the C-terminus of MyoA and TgELC2 or TgELC1 in both calcium-bound and -free forms as well as the X-ray crystal structure and NMR solution structures of the N-terminal domain of *P. falciparum* PfELC. We provide a thorough characterization of all identified interaction surfaces and demonstrate that the ELCs bind to a conserved binding site on MyoA. Furthermore, we show that the N-terminal domain of isolated PfELC is structured whereas its C-terminus is more disordered than its *T. gondii* homologs. However, ELCs from both *P. falciparum* and *T. gondii* mutually induce the structure with the disordered MyoA C-terminus to assemble into structurally conserved complexes.

## Results

### PfELC folds into a calmodulin-like N-terminal domain with a disordered C-terminus

Crystal structures of *T. gondii* and *P. falciparum* MyoA [23, 24] as well as structures of their distal light chains MLC1 (MTIP) [15, 22] have already been determined. To shed light on the architecture and folding of the recently identified proximal essential light chains (ELCs) we studied their structure in the context of their interaction partners. Analysis of the sequences of the two *T. gondii* myosin essential light chains TgELC1 and TgELC2 and the *P. falciparum* homolog PfELC indicates likely structural differences for the latter. TgELC1 and TgELC2 share a high degree of sequence identity (44.4%) and similarity (65.2%), whereas PfELC is significantly less similar to the *T. gondii* ELCs, with 20.3% identity and 40.6% similarity to TgELC1 (Fig 1C). Likewise, the disorder probability differs between the *T. gondii* and *P. falciparum* essential light chain homologs (S1A Fig). To study the structural differences, we recombinantly expressed N-terminally His-tagged ELCs in *E. coli* (Fig 1B) and purified them to homogeneity. In spite of similar molecular weight, PfELC and TgELC2 display distinct elution profiles when subjected to size-exclusion chromatography (SEC). PfELC elutes earlier compared to TgELC2 (Fig 2A), suggesting differences in the hydrodynamic radius of these constructs. Small angle X-ray scattering (SAXS) measurements further confirm that PfELC does indeed have a larger overall size in solution compared to that of TgELC2, with the respective radii of gyration (*R_g_*) being 2.71 ± 0.05 nm and 2.14 ± 0.05 nm (Fig S1B-D, Table 1 and S2 Table). The SAXS data also provide evidence that the increased *R_g_*of PfELC likely results from conformational flexibility, as the peak of PfELC in the dimensionless Kratky plot is broader and shifted towards higher angles compared to that of TgELC2 (Fig S1C, S2 Table). This finding is corroborated by the observed secondary structure content for PfELC derived from circular dichroism spectroscopy, showing that PfELC has lower α-helical and higher random coil content compared to TgELC2 (Fig 2B, Table 1). In order to map structured elements and disordered regions of PfELC, we performed triple-resonance NMR experiments that facilitated the near complete assignment of the amide backbone resonances (Fig. S1E). Secondary structure elements were determined from chemical shifts and the dynamics of the PfELC backbone was probed using heteronuclear NOEs ({^1^H}-^15^N NOE). This ^15^N based dynamics experiment allows to distinguish between rigid ({^1^H}-^15^N NOE > 0.7, secondary structure elements), somewhat flexible ({^1^H}-^15^N NOE ∼ 0.5-0.7, loops and turns) and extremely flexible ({^1^H}-^15^N NOE < 0.5, unfolded/ random coil) regions of the protein.

**Fig 2.**
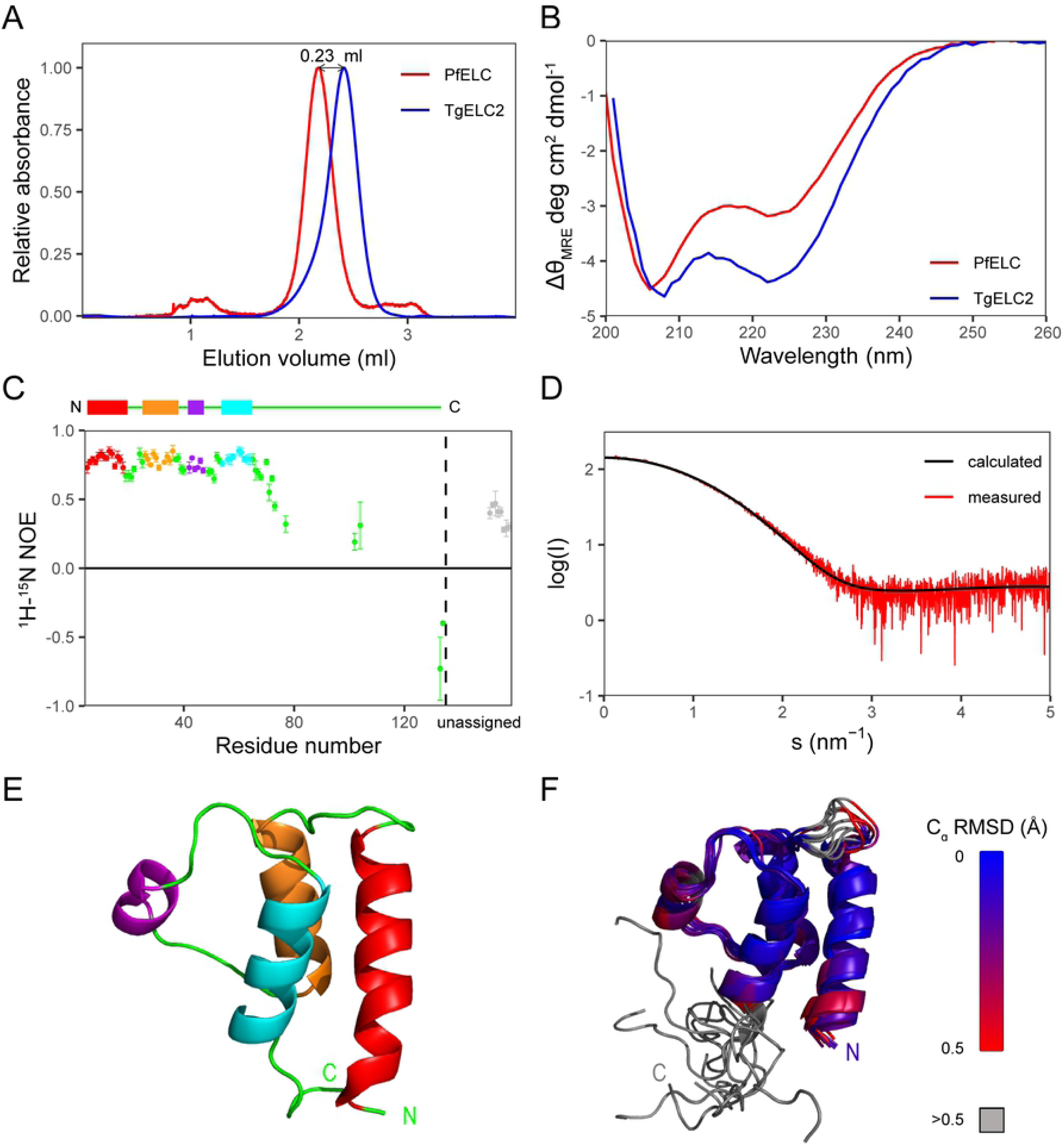
Structural differences between ELCs of *T. gondii* and *P. falciparum*. (A) Gel filtration profile of PfELC (red) and TgELC2 (blue) on a home-packed Superdex 200 5/150 column. PfELC elutes at a smaller elution volume, suggesting that it has a larger hydrodynamic radius compared to TgELC2. (B) Far-UV-circular dichroism spectrum of PfELC (red) and TgELC2 (blue) shows that PfELC has a lower α-helical and higher random coil content compared to TgELC2. (C) Backbone dynamics of PfELC on a picosecond to nanosecond time scale. Heteronuclear NOE ({^1^H}-^15^N NOE) of PfELC on a residue basis. Residues are colored according to secondary structure elements (four α-helices: from N terminus red, orange, violet, cyan, random coil/loop residues are green, unassigned C-terminal residues in grey). (D) Experimental small angle X ray scattering curve of PfELC (red) and calculated scattering (black line) from the crystal structure of the PfELC monomer fit with a Χ^2^ value of 1.37, confirming that the protein is a structurally rigid globular monomer in solution. (E) Crystal structure of the N-terminal isolated domain of PfELC, residues 1-68. PfELC displays a typical calmodulin fold with two helix-loop-helix motifs. The degenerate EF hand loops do not bind any ion. In agreement with NMR data of full length PfELC, the protein consists of four α-helices (from N terminus red, orange, violet, cyan, loops and disordered regions in green). The termini are labelled. (F) Ten lowest-energy NMR structures of PfELC (residues 1-74) colored from lowest (blue) to highest (red) backbone RMSD compared to the crystal structure show that the loop of the PfELC first EF hand (residues 16-22) and the third helix (residues 40-47) display a certain degree of flexibility.

**Table 1.** Biophysical characterization and comparison of PfELC and TgELC2.

|  | SEC data | CD data |  |  | SAXS data |  |  |
| --- | --- | --- | --- | --- | --- | --- | --- |
| | Elution volume (ml) | $\alpha$ helix | $\beta$ sheet | random | Rg (nm) | MW (kDa) | Dmax (nm) |
| **TgELC2** | 2.41 | 41% | 15% | 45% | 2.15 | 17.0 | 6.73 |
| **PfELC** | 2.18 | 34% | 16% | 51% | 2.83 | 16.3 | 9.50 |

This analysis on PfELC revealed that the protein contains an α-helical structure in the N-terminal domain, while the C-terminal part is disordered in isolation as evident by the low heteronuclear NOEs for this region (Fig 2C). Based on this finding, we expressed and purified an N-terminal fragment of PfELC (amino acids 1-74, PfELC-N; see Fig 1B) and determined its structure by both X-ray crystallography to 1.5 Å resolution (Fig 2E, Table 2) and by NMR spectroscopy (Fig 2F, S3 Table). The ten lowest energy NMR structures superimpose with an average backbone RMSD of 0.87 Å and an all-atom RMSD of 1.23 Å. The lowest energy NMR conformer is very similar to the crystal structure, with a backbone RMSD of 1.4 Å over residues 1-68. The N-terminal domain of PfELC has a typical calmodulin fold with two EF-hands formed by two helix-loop-helix motifs. EF-hands typically have the capacity to bind calcium [25] but here, both EF-hands lack the canonical calcium binding residues and therefore are not able to bind calcium as evident in the determined crystal structure. PfELC-N crystallized as a dimer with a covalently linked disulfide bridge between cysteine 19 residues on both protein chains (S1F Fig). However, in solution, the protein is monomeric, as shown on non-reducing SDS-PAGE (S1G Fig) and by SAXS (the scattering computed from the crystal structure yields a good fit with discrepancy Χ^2^=1.37 to the SAXS data, Fig 2D, S2 Table), and cysteine 19 is reduced, as probed by the indicative NMR ^13^Cβ shift. A comparison of the crystal structure with the NMR structure highlights that the loop of the first EF hand (residues 16-22) and the third helix (residues 40-47) displays the highest degree of flexibility, in agreement with the heteronuclear NOE experiment (Fig 2C), while the position of the other loops agrees well between the crystal and the NMR structure (Fig 2E-F). In general, the assigned backbone resonances in the NMR spectra superimpose for both full-length protein PfELC and the N-terminal domain, highlighting that the N-terminal domain maintains the same structure in both constructs (S1E Fig). These results show that isolated PfELC is monomeric in solution with a calmodulin-like N-terminal fold and a disordered C-terminal region.

**Table 2.**
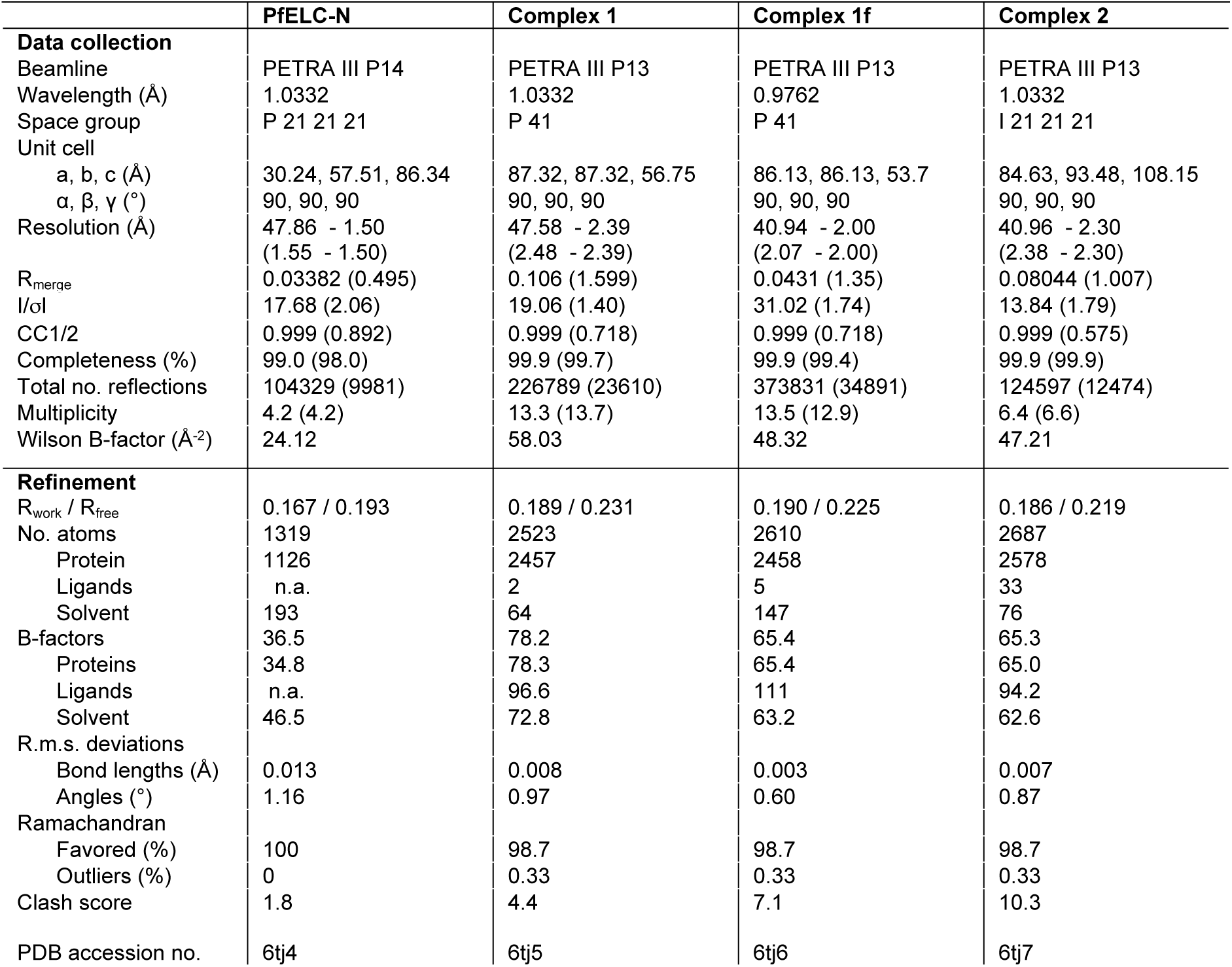
X-ray data collection and refinement statistics.

### Essential light chains bind conserved sequence of MyoA

Both *T. gondii* ELCs (TgELC1 and TgELC2) as well as *P. falciparum* PfELC have been previously shown to bind to the C-terminus of MyoA [13,14,17]. Whereas for PfELC, two binding sites of the PfMyoA C-terminus (PfMyoA residues 786-803 and 801-818; Fig 3A) were identified [14], only one binding site was experimentally confirmed for TgELCs (TgMyoA 775-795; Fig 3A) [15, 16]. To investigate these interactions further, we measured the binding affinity of TgELC1, TgELC2 and PfELC to peptides corresponding to the proposed MyoA binding sites.

**Fig 3.**
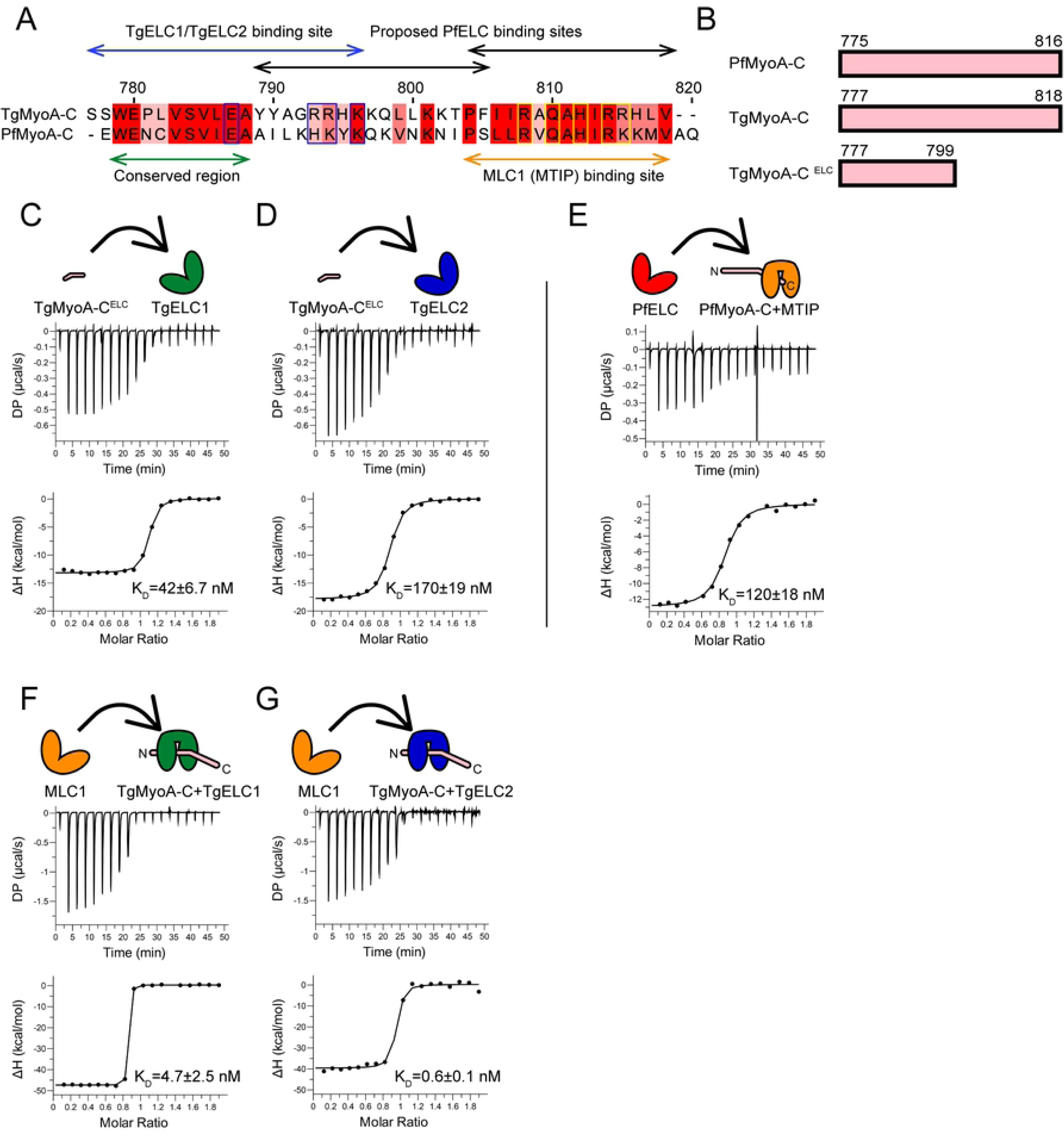
Formation of trimeric complexes between ELC, MLC1 (MTIP) and the MyoA C-terminus. (A) Sequence comparison of TgMyoA and PfMyoA C-termini shows a conserved region (green arrow) upstream of the MLC1 (MTIP) binding site. Whereas two binding sites of PfELC at the very C-terminus of PfMyoA were proposed (black arrows) [14], our data show that the actual binding site of PfELC encompasses the MyoA conserved region and is similar to the TgELC/TgMyoA binding site (blue arrows). The blue boxed residues indicate residues involved in polar interactions with TgELC1 and TgELC2, while yellow boxed residues form polar interactions with MLC1 (see Fig 5C-D and S4 Table). (B) The peptide constructs representing the C-terminal regions of MyoAs with indicated domain boarders used in this study. The constructs PfMyoA-C and TgMyoA-C encompass both MLC1/MTIP binding sites as well as the upstream conserved region which binds the essential light chains. The construct TgMyoA-C^ELC^ only consists of the TgELC binding site. (C,D) Isothermal titration of TgMyoA-C^ELC^ with TgELC1 and TgELC2 show that both dimeric complexes form with nanomolar affinity. The upper panel shows the signal recorded directly after each injection of TgELC1 and TgELC2 and represents the thermal power that has to be applied to maintain a constant temperature in the sample cell during recurring injections. In the lower panel, the integrated heats are plotted against the peptide/protein concentration ratio. The thermodynamic binding parameters were obtained by nonlinear regression of the experimental data using a one-site binding model. (E) Binding isotherm of PfELC titrated to the preformed MTIP/PfMyoA-C complex proves that the conserved hydrophobic region of MyoAs is indispensable for ELC binding. (G,H) Binding isotherms of MLC1 titrated into the pre-complex of TgMyoA-C with TgELC1 and TgELC2. MLC1 binds the pre-complex with high nanomolar affinity. All thermodynamic parameters derived from ITC measurements are summarized in Table 3.

**Table 3.** Overview of thermodynamic constants measured by ITC.

| **Dimeric interactions** |  |  |  |  |  |
| --- | --- | --- | --- | --- | --- |
| Protein (cell) | MyoA peptide (syringe) | Molar ratio | K<sub>d</sub> (nM) | ΔH (kcal/mol) | -TΔS (kcal/mol) |
| MTIP | PfMyoA-C<sup>ELC</sup> | 0.71 ± 0.01 | 320 ± 57 | -15 ± 0.4 | 5.5 |
| TgELC1 | TgMyoA-C<sup>ELC</sup> | 1.05 ± 0.01 | 42 ± 6.7 | -13 ± 0.2 | 3.2 |
| TgELC2 | TgMyoA-C<sup>ELC</sup> | 0.87 ± 0.01 | 160 ± 19 | -17 ± 0.2 | 7.4 |
| TgELC2<sup>E10A</sup> | TgMyoA-C<sup>ELC</sup> | 0.79 ± 0.01 | 190 ± 25 | -17 ± 0.3 | 8.2 |
| TgELC2<sup>D16A</sup> | TgMyoA-C<sup>ELC</sup> | 0.80 ± 0.01 | 280 ± 30 | -19 ± 0.3 | 9.7 |
| TgELC2<sup>F79A</sup> | TgMyoA-C<sup>ELC</sup> | 0.84 ± 0.01 | 280 ± 34 | -18 ± 0.3 | 9.5 |
| TgELC2<sup>S101A</sup> | TgMyoA-C<sup>ELC</sup> | 0.88 ± 0.02 | 280 ± 85 | -18 ± 0.8 | 9.3 |
| TgELC2<sup>E10A+H110A</sup> | TgMyoA-C<sup>ELC</sup> | 0.75 ± 0.02 | 1100 ± 220 | -21 ± 0.9 | 12 |

| **Trimeric interactions** |  |  |  |  |  |
| --- | --- | --- | --- | --- | --- |
| Pre-complex with MyoA-C (cell) | Protein (syringe) | Molar ratio | K<sub>d</sub> (nM) | ΔH (kcal/mol) | -TΔS (kcal/mol) |
| MTIP | PfELC | 0.83 ± 0.01 | 120 ± 18 | -13.0 ± 0.25 | 3.6 |
| TgELC1 | MLC1 | 0.92 ± 0.01 | 4.7 ± 2.5 | -39.1 ± 0.76 | 28 |
| TgELC2 | MLC1 | 0.81 ± 0.01 | 0.6 ± 0.1 | -47.6 ± 0.12 | 35 |
| TgELC2<sup>R17A</sup> | MLC1 | 0.92 ± 0.01 | 4.6 ± 0.4 | -49.7 ± 0.17 | 38 |
| TgELC2<sup>E22A</sup> | MLC1 | 0.92 ± 0.01 | 5.2 ± 1.9 | -45.9 ± 0.65 | 35 |
| TgELC2 | MLC1<sup>K168A</sup> | 0.79 ± 0.01 | 1.2 ± 0.8 | -47.7 ± 0.24 | 36 |
| TgELC2 | MLC1<sup>Q169A</sup> | 0.89 ± 0.01 | 2.3 ± 1.9 | -48.8 ± 0.50 | 37 |
| TgELC2 | MLC1<sup>N172A</sup> | 0.84 ± 0.01 | 4.3 ± 4.3 | -41.6 ± 0.92 | 30 |

The thermodynamic parameters were fitted by a one site binding model with the MicroCal PEAQ-ITC Analysis Software.

Strikingly, we could not monitor any binding of PfELC to the previously described binding sites but observed precipitation upon mixing PfELC with the respective peptides. The TgELC1/2 binding site is highly conserved between *T. gondii* TgMyoA and *P. falciparum* PfMyoA, therefore we hypothesized that PfELC could bind the homologous conserved region (see Fig 3A) and extended the PfMyoA peptide accordingly. However, precipitation occurred again and we speculated that the presence of MTIP bound to PfMyoA is a prerequisite for binding of PfELC. In agreement with this hypothesis, previous reports have shown that PfELC can be co-purified with full-length MyoA only in the presence of MTIP from insect cells [17]. To determine the affinity of PfELC via isothermal titration calorimetry we first formed a complex between MTIP and the PfMyoA neck region peptide (PfMyoA-C, residues 775-816; S2 Fig and see Fig 3B) and then titrated in PfELC. PfELC bound to MTIP and PfMyoA-C with an affinity of 120 ± 18 nM (Fig 3E and Table 3). Interestingly, we observed the opposite behavior for the *T. gondii* light chains. The *T. gondii* MTIP homolog, MLC1, precipitated upon interacting with TgMyoA-C (residues 777-818, see Fig 3B), but both TgELC1 and TgELC2 bound TgMyoA-C^ELC^ (residues 777-799, see Fig 3B) with high affinity (42 ± 6.7 nM and 160 ± 19 nM, respectively) (Fig 3C-D and Table 3).

Finally, we were able to reconstitute the trimeric complex with both TgELC1 and TgELC2, by first forming the dimeric complex between TgELC1 or TgELC2 and TgMyoA-C and then adding MLC1. MLC1 bound the preformed complex of TgELC1 or TgELC2 and TgMyoA-C with high affinity (4.7 ± 2.5 nM and 0.6 ± 0.1 nM, respectively) (Fig F-G and Table 3). In conclusion, *T. gondii* TgELC1 and TgELC2 need to interact with TgMyoA-C *in vitro* first and only then MLC1 can assemble into the trimeric complex. In *P. falciparum*, MTIP first interacts with the C-terminus of PfMyoA and only after that, PfELC is able to bind and form the trimeric complex. All ELC homologs bind a highly conserved sequence stretch at the C-terminus of MyoA (Fig 3A).

### Essential light chains bind MyoA in a compact conformation and induce an α-helical structure

Previous reports have shown that the presence of *P. falciparum* and *T. gondii* essential light chains increases the speed of the myosin A motor twofold [14,16,17]. To understand the functional role of ELCs on a molecular level, we characterized TgELC2 in a free and bound state in complex with TgMyoA-C^ELC^ (see Fig 3B). On size exclusion chromatography, the dimeric complex of TgELC2 and TgMyoA-C^ELC^ elutes later than TgELC2 alone, indicating that the overall size of TgELC2 decreases upon binding of TgMyoA-C^ELC^ (Fig 4A). Indeed, the parameters extracted from the SAXS measurements indicate compaction upon formation of the complex (see S2 Table and Fig 4B-C). Before addition of the MyoA-C^ELC^, the flexible nature of TgELC2 relative to the complex is observed as a broad peak shifted to high angles in the dimensionless Kratky plot. Upon complex formation, the peak becomes narrower and shifts to lower angles, an indication of macromolecular compaction (Fig 4B, also see S3A-B Fig). In agreement, the *R_g_* calculated from the experimental data by Guinier analysis [26] decreases from 2.15 nm to 1.73 nm upon binding and the particle distance distribution changes accordingly, with the maximum size decreasing from 6.7 nm for TgELC2 to 5.5 nm for the complex (S2 Table and Fig 4C). These data suggest that the flexible TgELC2 protein undergoes a conformational change upon binding to TgMyoA C-terminus, adopting a compact structure, similarly to what was reported for the MTIP-PfMyoA interaction [22]. In all previously solved myosin structures, the myosin neck regions fold in a long α helix, tightly bound by their light chains [27]. This rigid conformation allows the neck region to act as the lever arm of myosin and its stiffness directly correlates with the myosin step size and speed [28–30]. However, both TgMyoA-C as well as PfMyoA-C are unfolded or partially unfolded in isolation (S3C Fig). Indeed, the C-terminal amino acid residues of the recently published TgMyoA [23] and PfMyoA [24] motor domain structures could not be resolved, likely due to their intrinsically disordered nature. We hypothesized that the essential light chains can induce α-helical structure in MyoA upon binding, therefore we measured circular dichroism of TgMyoA-C^ELC^ and TgELC2 in isolation and in the complex (Fig 4D). The data show that isolated TgMyoA-C^ELC^ is unstructured and TgELC2 has a predominantly α-helical fold. However, the circular dichroism spectrum of the dimeric complex has a significantly higher α-helical content than the sum of the spectra of the two individual components, as shown by a lower ellipticity at 222 nm and a higher ellipticity at 195 nm, suggesting that the content of the α-helical secondary structure increased upon formation of the complex. We observed similar, albeit less pronounced effect also for the TgELC1-TgMyoA-C^ELC^ complex assembly (S3D Fig). We anticipate that the increase in α-helical secondary structure content corresponds to the induction of the structure of the TgMyoA C-terminus, which in turn stiffens the TgMyoA lever arm and enhances the performance of TgMyoA in the full-length context.

**Fig 4.**
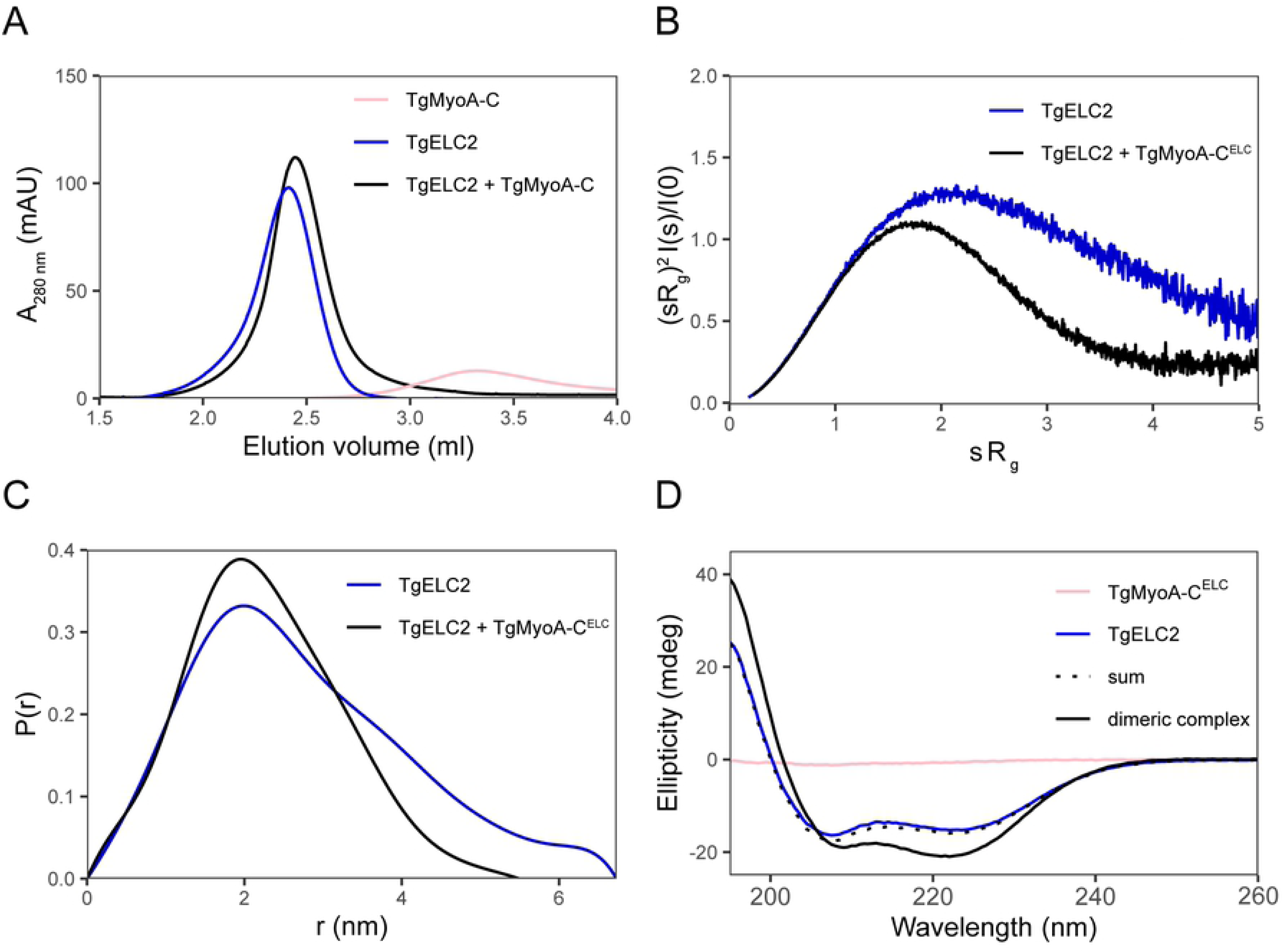
Both TgELCs and TgMyoA undergo large conformational changes upon binding. (A) Dimeric complex of TgELC2 and TgMyoA-C elutes at shorter retention times than isolated TgELC2 on a home-packed Superdex 200 5/150 column, suggesting that the hydrodynamic radius of TgELC2 decreases upon TgMyoA-C binding. (B) Kratky plots of isolated TgELC2 and in complex with TgMyoA-C. The dimensionless Kratky plot of TgELC2 in complex with TgMyoA-C^ELC^ (black) has a maximum close to *sR_g_ =* √3 and converges to zero, unlike isolated TgELC2 (blue), suggesting that TgELC2 in isolation is rather extended and compacts upon binding to TgMyoA. (C) The distance distribution calculated by Guinier analysis from the SAXS data further confirms that TgELC2 undergoes compaction upon TgMyoA binding. TgELC2 displays wider distance distribution with *d_max_* = 6.7 nm, whereas the distance distribution of the dimeric complex is narrower with *d_max_* = 5.5 nm. (D) The far-UV CD data indicate that TgELC2 induces a α-helical structure in TgMyoA upon binding. The individual spectra of TgELC2 and TgMyoA-C^ELC^ do not sum up to the CD spectrum of their dimeric complex and the CD spectrum of the dimeric complex displays more pronounced features of α-helical secondary structure with lower ellipticity at 222 nm and higher ellipticity at 195 nm compared to the sum of individual components. CD spectra were recorded in a 1 mm cuvette at a concentration of 5 μM of each component in 10 mM NaP (pH 7.5), 150 mM NaF and 0.25 mM TCEP at 20°C.

### TgELC1 and TgELC2 form structurally similar complexes with MyoA and MLC1

To gain detailed information on the architecture of the trimeric protein assemblies, we crystallized and determined the crystal structures of the following trimeric complexes: *T. gondii* MLC1/TgMyoA-C/TgELC1 complex at 2.4 Å resolution (hereafter named complex 1) and *T. gondii* MLC1/TgMyoA-C/TgELC2 complex at 2.3 Å resolution (hereafter named complex 2) (Fig 5A-B, Table 2). Overall, both complexes display a similar architecture. MyoA folds into an extended α helix with a characteristic kink between residues 801-803 (angle of 139° in complex 1 and 137° in complex 2). Both TgELCs display a typical calmodulin fold with one N-terminal and one C-terminal lobe, each lobe comprised of two EF hands. With the exception of the first EF hand, all EF hand sequences deviate from the canonical EF hand sequence and as expected, do not bind any ions. However, clear additional electron density was visible for the first EF hand and assigned to a bound calcium ion coordinated in a tetragonal-bipyramidal geometry. Both TgELCs form conserved polar interactions with TgMyoA, involving TgMyoA residues E787, R793, R794 and K796, a π-π stacking interaction between the conserved residue pair W779-F79 and a group of hydrophobic residues clustered around the conserved TgMyoA region P801-Y810 (Fig 5C-D, S4 Table). Mutational analysis on TgELC2 (Table 3, S4A Fig) showed that disrupting one of the polar interactions or the conserved π-π stacking interaction W779-F77 only has a minor effect on the binding affinity of TgMyoA to TgELC2 and suggests that the hydrophobic residues in the conserved MyoA region play a crucial role for complex formation. In general, TgELC1 forms tighter interactions with TgMyoA-C than TgELC2, with a higher number of interatomic interactions and a larger protein-protein interface (S4 table, Fig 5C-D), which is consistent with the difference in the binding affinity of the dimeric complexes measured by ITC (Fig 3C-D, Table 3).

**Fig 5.**
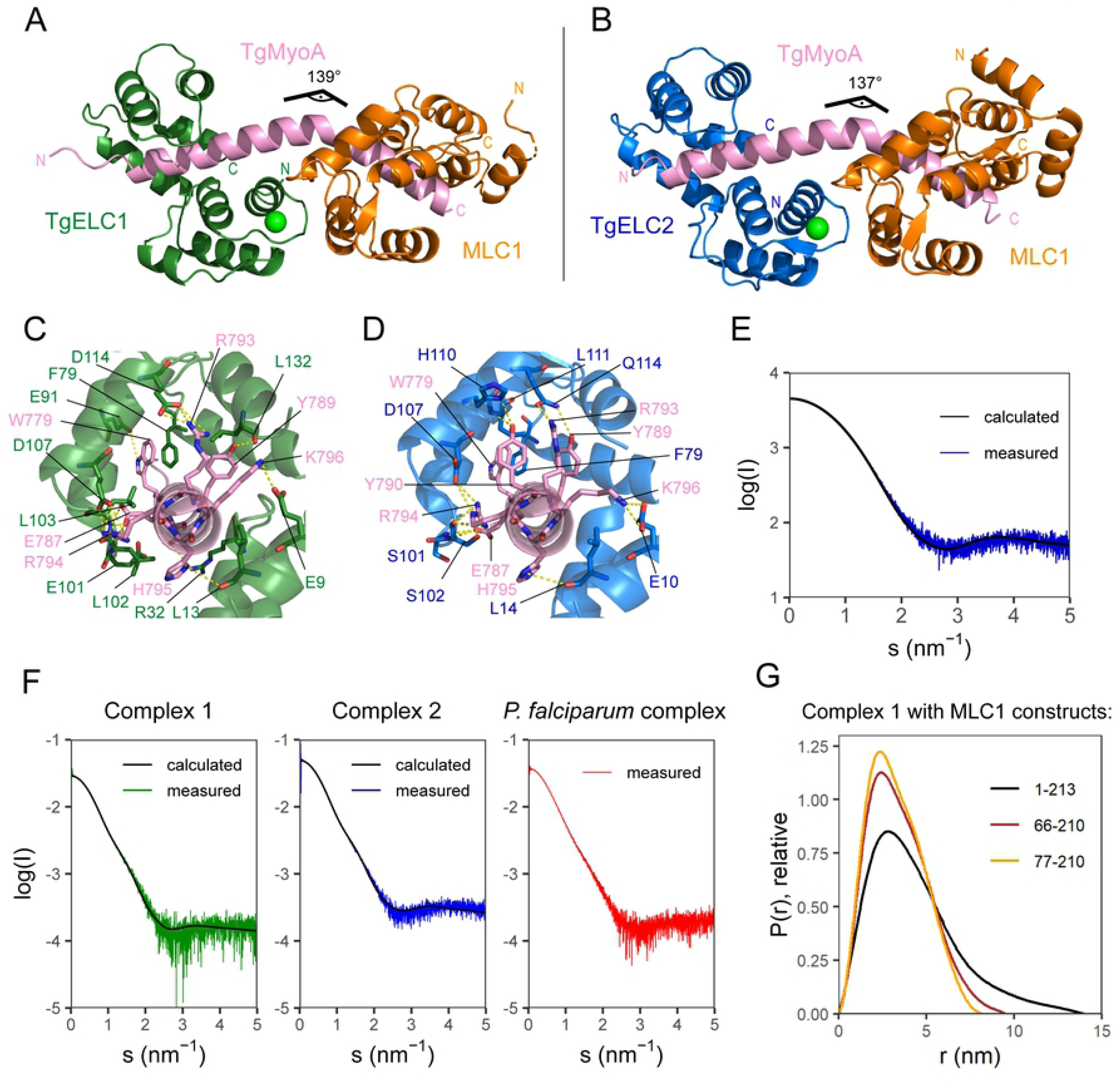
Crystal structures of glideosome trimeric complexes. (A) Crystal structures of trimeric complex of TgELC1 (green), TgMyoA-C (pink) and MLC1 (orange) (complex 1). (B) Crystal structure of trimeric complex of TgELC2 (blue), TgMyoA-C (pink) and MLC1 (orange) (complex 2). The complex structures with TgELC1 and TgELC2 are very similar and the TgMyoA helix displays a characteristic kink between residues 801-803. ELCs bind upstream of the MLC1 binding site. (C,D) Binding interface between TgMyoA-C (pink) and TgELC1 (green) in complex 1 and TgELC2 (blue) in complex 2. Residues involved in polar interactions are labelled with the corresponding colour and shown in stick representation. Most polar interactions are mediated by C-terminal lobes of TgELCs and the hydrophobic interactions between TgELCs and the conserved hydrophobic TgMyoA residues play a crucial role in complex formation as evident from ITC measurements. (E) Experimental small angle X-ray scattering curve of TgELC2 bound to TgMyoA-C^ELC^ (blue). The fit to the scattering pattern computed from the crystal structure of complex 2 omitting MLC1 (black line, χ^2^ = 1.16) shows that TgELC2 does not undergo major conformational changes upon binding of MLC1 and the formation of the trimeric complex. (F) SAXS analysis of the different trimeric complexes. Similarity of the scattering profiles indicates a similar shape for all three complexes, suggesting that the architecture of the trimeric complexes is conserved between *T. gondii* and *P. falciparum*. Computed scattering curves of complex 1 and complex 2 fit the respective experimental data with χ^2^ = 1.26 and χ^2^ = 2.41, respectively. (G) The distance distribution plots of complex 1 calculated from experimental small angle X ray scattering data change by shortening the MLC1 N-terminal domain, indicating flexibility of MLC1 upstream of residue 77. The distance distribution is narrower upon N-terminal truncation of MLC1, with *d_max_* decreasing from 14 nm (complex 1 with full-length MLC1^1-213^) to 9.5 nm (MLC1^66-210^) and further to 8.2 nm (MLC1^77-210^).

This tighter interaction is also reflected in the SAXS data measured for both trimeric complexes where the extracted molecular weight estimates are consistent with the expected molecular weights (Fig 5F, S2 Table). Complexes 1 and 2 are monomeric in solution but whereas the calculated scattering data of complex 1 fit the experimental scattering data with a Χ^2^ of 1.26, the structure of complex 2 displays a higher Χ^2^ = 2.41, suggesting some structural differences in solution. In addition, the scattering data of the *P. falciparum* trimeric complex of MTIP, PfELC and PfMyoA-C are very similar to the *T. gondii* homologues, suggesting that the overall architecture of the complexes is conserved between *P. falciparum* and *T. gondii* (Fig 5F, S2 Table).

To investigate whether MLC1 binding to the preformed dimeric complex induces additional conformational changes in TgELC2, we recorded SAXS data of the TgELC2-TgMyoA-C^ELC^ dimeric complex (Fig 5E). The resulting scattering data are in excellent agreement with the calculated scattering profile from the trimeric complex omitting MLC1 (Χ^2^ of 1.16 Å) indicating no major structural rearrangements. Similarly, MLC1 adopts the same conformation as in the already described structure of a MLC1-MyoA^803-830^ dimeric complex (PDB ID 5vt9), with a backbone RMSD of 0.96 Å compared to complex 1 and 0.75 Å compared to complex 2 (S4B-C Fig). The key interactions of TgMyoA with MLC1 are also conserved between the structures of the trimeric and dimeric complexes (R808, H812, R814) with the exception of few weak polar interactions (see S4 Table).

In general, the N-termini of MLC1 and its *P. falciparum* homolog MTIP were shown to anchor myosin A to the inner membrane complex by interacting with GAP45 *in vivo* [31]. Interestingly, although the same construct of MLC1 was used for crystallization in all cases (residues 66-210), the quality of the electron density only allowed to build the N-terminal part of the model to different extents in complex 1 (from residue 77), complex 2 (from residue 67) and the previously solved structure of the dimeric complex of MLC1 and TgMyoA C-terminus (from residues 73, 5vt9, [15]). These differences indicate that the N-terminal residues of MLC1^66-210^ in solution show a certain degree of disorder. To further investigate the structure of the N-terminal residues of MLC1 in solution, we recorded SAXS data of complex 1 with full-length MLC1 (residues 1-213) and two MLC1 constructs with a truncated N-terminus (MLC1^66-210^, which was also used for crystallization, and MLC1^77-210^; Fig 5F-G, S4D Fig).

The complex containing full-length MLC1 displays a significantly higher maximum particle size (D_max_=14 nm) and a larger radius of gyration (3.50±0.02 nm) in comparison to the complex used for crystallization (S2 Table), suggesting that the MLC1 N-terminus is disordered in the complex with TgMyoA and TgELC1. The D_max_ decreases even further from 9.5 nm to 8.2 nm upon truncating the MLC1 construct by additional eleven residues (from MLC1^66-210^ to MLC1^77-210^). However, the eleven N-terminal residues (66-76) of MLC1 in complex 2 form an α helix that folds back towards the center of the molecule and therefore does not effectively increase the maximum particle size of the molecule. The difference in the measured particle size thus indicates that the structural elements upstream of residue 77 of MLC1 are flexible in solution (Fig 5G). Moreover, the SAXS data of complex 1 with the shortest MLC1 construct (77-210) agree well with the calculated scattering profile of the crystal structure (Χ^2^=1.04), whereas the fit of the complex with MLC1^66-210^ is slightly poorer (Χ^2^=1.26), further supporting that residues 66-76, which could not be resolved in all crystal structures, are at least partially disordered. The flexibility within residues 66-76 is additionally apparent from the normal mode analysis (S6A Fig, see below). Thus, MLC1 residues 77-213 form a rigid complex with the C-terminus of TgMyoA, whereas residues 1-76 are flexible. This feature may have further implications on the function of the protein, namely anchoring MyoA to the membranes of the IMC or interacting with other members of glideosome, such as GAP45.

### Calcium stabilizes the trimeric complexes by mediating ELC interactions to MLC1

Apicomplexan invasion is a tightly regulated process, which also involves an increase in intracellular calcium concentration. To investigate the role of calcium bound in the first EF hand of both TgELCs, we determined an additional crystal structure of the calcium-free complex TgELC1/MLC1/MyoA-C at 2.0 Å (complex 1f, Fig 6A, Table 2). The calcium-free complex generally adopts the same conformation as complex 1. The MyoA-C helix is kinked at a similar angle (134°), and the binding interfaces between MLC1 and TgMyoA as well as between TgELC1 and MyoA are identical to complex 1 (S4 Table). The calcium binding residues remain in the same conformation as in complex 1 except for side chain of aspartate 17 which is flipped by 120 degrees and thereby enables the release of calcium from the binding pocket (Fig 6B). In complex 1, calcium is coordinated in a tetragonal bipyramidal geometry by the carboxyl groups of side chains D15, D17, D19, the carbonyl group of E21 and two water molecules. In complex 2, calcium is similarly coordinated by the homologous side chain residues of D16, N18, D20, the carbonyl group of E22 and two water molecules. Additionally, in complex 2, these water molecules are further stabilized by interactions with the side chains of E27 and Q49. Contrary to the *T. gondii* TgELCs, the homologous EF hand loop of PfELC is bent to the other side and does not possess the residues needed for coordination of calcium (Fig 6B). In agreement with the presented crystal structures, calcium has no major influence on the secondary structure of individual TgELCs or PfELC in solution (S5A-C Fig). Powell et al. recently showed that the absence of calcium reduces the affinity of TgELC1 for the MyoA C-terminus [15]. To investigate this effect in TgELC2, we mutated the crucial calcium binding residue D16 in the first EF hand to alanine. As expected, the complex of TgELC2^D16A^/MyoA-C^ELC^ could not be stabilized against heat unfolding by the addition of calcium (Fig 6C). However, the complex of TgELC2^D16A^/MyoA-C^ELC^ was also less stable compared to the wild type complex independent of the bound calcium and the effect of the mutation on the affinity of the dimeric complex was negligible (Table 3 and S5B Fig). Based on the available crystal structure we speculated that calcium ions might be crucial to form an interaction surface between the EF hand of TgELCs and MLC1. The structures of the trimeric complexes show several polar interactions in this interface (6D-F Fig, S4 Table). Therefore, we mutated selected TgELC2 or MLC1 residues at this interface to alanine and measured the binding affinity for trimeric complex formation. The observed decrease in affinity was only moderate (up to five-fold) but the measured affinities reached the limitations of reliable high affinity ITC measurements (Table 3 and S5C Fig). Therefore, we turned to stability measurements and observed a pronounced concentration dependent effect of calcium ions on the thermal stability of the entire complex in a concentration dependent manner (Fig 6G-H, S5D Fig). In conclusion, calcium binding by the first EF hand of TgELCs does not impact the formation of the complex but contributes substantially to the stability of the complex by maintaining the interface between MLC1 and TgELCs.

**Fig 6.**
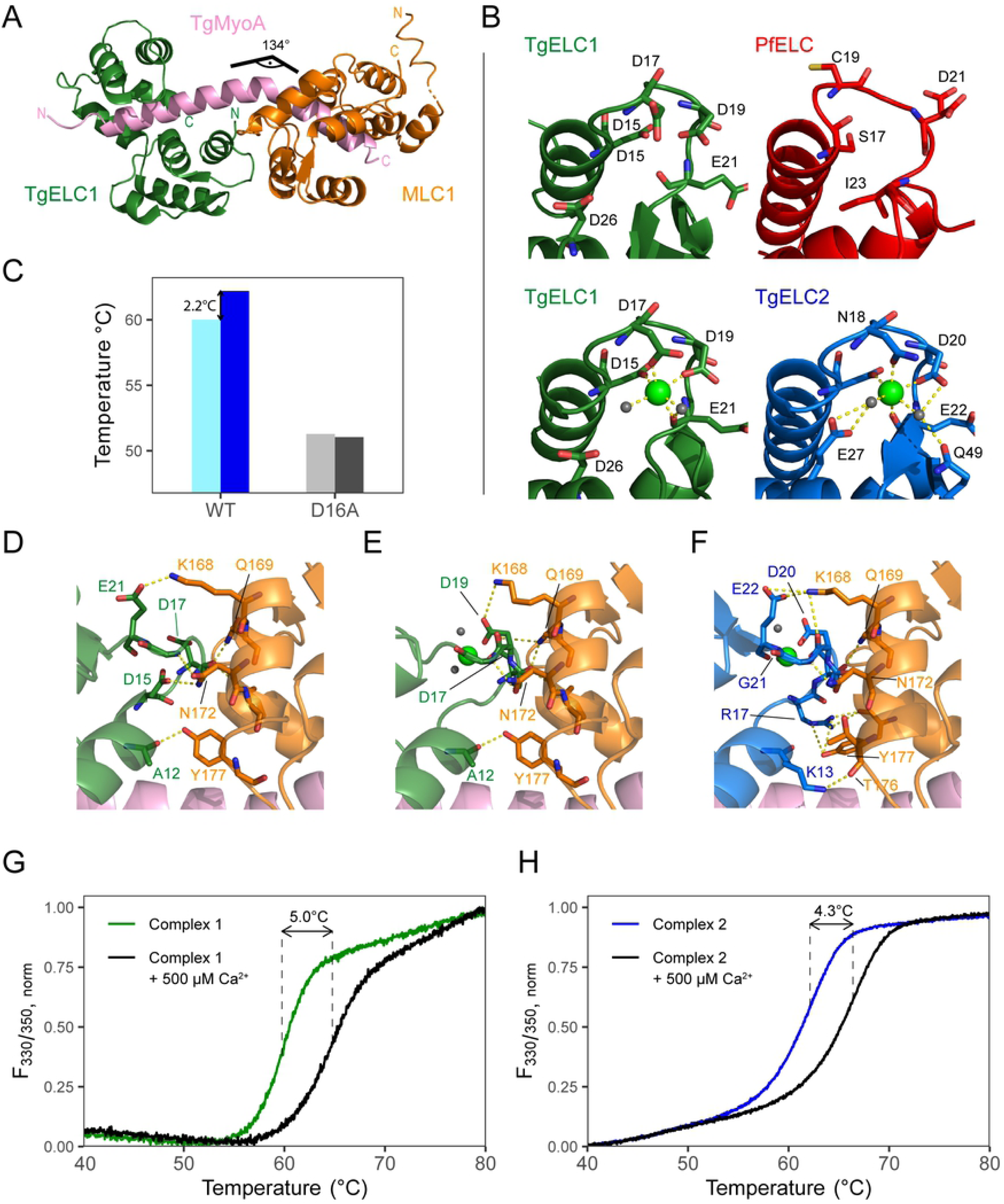
Influence of calcium on complex formation. (A) Crystal structure of the glideosome trimeric complex composed of TgELC1 (green), MLC1 (orange) and TgMyoA-C (pink) in the absence of calcium (complex 1f). The absence of calcium does not cause a major structural rearrangement (see Fig 5A). (B) Structural comparison of the first EF hand in ELCs and calcium coordination between complex 1f, complex 1, complex 2 and PfELC-N. Whereas PfELC does not bind any ion due to a degenerate sequence in its EF hand, both TgELCs in complex 1 and complex 2 bind calcium in a tetragonal bipyramid coordination, including two water molecules. These water molecules are further stabilized in complex 2 by additional residues (E27, Q49, D20). In complex 1f, the side chain of residue D17 is flipped by 120°, enabling the release of calcium. (C) Thermal stability of the dimeric complex of TgMyoA-C and wild type TgELC2 or its mutant TgELC2^D16A^ shows that the mutant TgELC2^D16A^ loses its ability to bind calcium but is also substantially less stable. (D-F) Binding interfaces between TgELCs and MLC1 in the trimeric complex structures of (from left) complex 1f, complex 1 and complex 2. Corresponding residues are labelled with the respective colour. The same set of residues (K168, Q169, N172, Y177) is involved in polar interactions (indicated by yellow dashes) on the MLC1 site, but various residues are utilized by TgELCs. (G-H) Thermal stability change of trimeric complex 1 and complex 2 upon addition of calcium measured by nanoDSF. The stability of both complexes strongly increases upon calcium binding.

### Trimeric complexes with full-length myosin A resemble the dynamics of conventional myosins in the pre-power stroke state

Previously solved structures of myosins in complex with their light chains suggest that the converter domains interact with the essential light chain to further stabilize the rigid lever arm and possibly transmit the structural changes from the myosin motor domain to the lever arm [32, 33]. Similar to that, it has been proposed that TgELC1 might constitute a small binding interface with the TgMyoA converter domain [15], providing enhanced rigidity to the myosin lever arm. To investigate whether the crystal structures of complex 1 and complex 2 are compatible with these observations and ensure that they do not clash with the TgMyoA core, we built structural models of the TgMyoA motor and neck domain bound to MLC1 and TgELC1 or TgELC2. For model building, we made use of the crystal structure of the TgMyoA motor domain in the pre-power stroke state (PDB ID 6due [23], residues 33-771) and extended its C-terminus by an α helix (TgMyoA residues 772-791), which resulted in 50 MyoA models. Subsequently, the MyoA residues 780-791 of the crystal structures of complex 1 or complex 2 were aligned to these models and five models of each complex with the lowest clash score were energy minimized. The details of model building are described in the Methods section. In all cases, the energy-minimized models did not contain any clashes, indicating that our structures are compatible within the full-length context of TgMyoA (Fig 7A-B). TgMyoA residues 762-818 constituting the lever arm maintained a continuous α helix after energy minimization with both TgELC1 and TgELC2 forming a small number of contacts with the TgMyoA converter domain. These contacts mainly involve the side chain of arginine 81 of TgELC1 or TgELC2 and residues 720-724 of TgMyoA, which is in agreement with the previously published HDX data [15]. To further explore the dynamics of full-length TgMyoA with its light chains, we performed normal mode analysis in an all-atom representation on five energy-minimized models from each complex 1 and complex 2, and subsequent deformation analysis which allowed us to identify potential hinge regions within these structures. In both cases, all five reconstructed models displayed nearly identical pattern of motions (see S6A Fig for complex 2): the structures undergo bending in the hinge region of TgMyoA residues 773-777 in two perpendicular directions (mode 7 and 8) as well as twisting in the same region (mode 9). In the remaining modes (modes 9 and higher), the movement further propagates throughout the lever arm helix up to TgMyoA residue 799. As a result, the deformation analysis of the 20 lowest energy modes predicted the hinge region of the TgMyoA lever arm between TgELCs and the converter domain, and an additional hinge between TgELCs and MLC1 (complex 2 in Fig 7C and complex 1 in S6B Fig). Such dynamics of the myosin light chains is similar as previously described in conventional myosins [33, 34] and the flexibility in the first TgMyoA hinge can contribute to the efficient rebinding of the myosin motor domain to actin in the pre-power stroke state (6SC Fig) [35]. In conclusion, the structures of the trimeric complexes composed of the TgMyoA light chains and TgMyoA C-terminus are compatible with the full-length TgMyoA and exhibit dynamics that is similar to the dynamics of conventional myosins.

**Fig 7.**
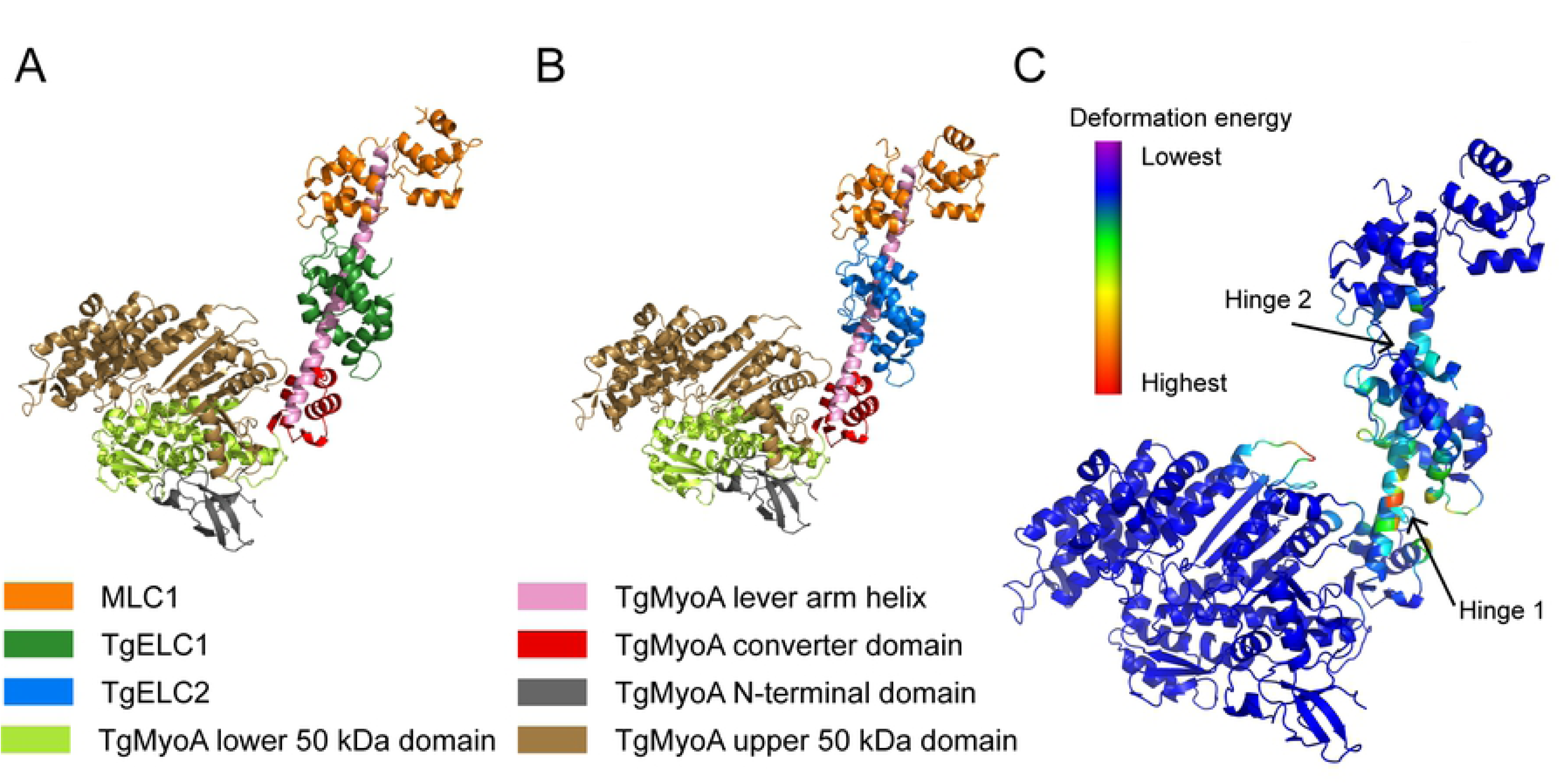
Trimeric complexes modelled in the full-length MyoA context. (A) Energy-minimized model of complex 1 as a part of TgMyoA. (B) Energy-minimized model of complex 2 as a part of TgMyoA. The models show that the crystal structures of the trimeric complexes are compatible with the structure of TgMyoA and maintain the α-helical structure of the TgMyoA lever arm. No clashes between TgMyoA and TgELCs were observed. (C) Deformation analysis of complex 2 identified two hinge regions in the lever arm of myosin A, which contribute to most of the observed dynamics of the protein complex within the 20 lowest-energy modes. The model is coloured by deformation energy from lowest (violet) to highest (red). The hinges localize to the TgMyoA lever arm between the converter domain and the TgELC2 binding site (hinge 1, residues 773-777) as well as between the TgELC2 and MLC1 binding sites (hinge 2, residues 799-801). These deformations agree with the role of TgMyoA in the pre-power stroke state in the context of a power stroke cycle, where the myosin is probing the conformational space to bind to actin.

## Discussion

The gliding motility of apicomplexan parasites is generated by a myosin-A-based molecular motor that is anchored in the inner apicomplexan membranes by a multi-protein complex called the glideosome. While the structures of some individual glideosome components and small dimeric complexes have been solved for *T. gondii* and *P. falciparum* so far, they do not explain their roles in the glideosome assembly and regulation. In this study, we present structures of trimeric glideosome complexes consisting of the TgMyoA C-terminus, MLC1 and TgELC1 or TgELC2 and investigate the role of essential light chains in glideosome assembly and function.

### Binding of essential light chains and induced folding

We showed that the α-helical content of *P. falciparum* PfELC is lower compared to *T. gondii* ELCs and that the C-terminus of PfELC in isolation is disordered. However, the α-helical secondary structure content of *T. gondii* ELCs is even higher in the above described crystal structures of the trimeric complexes compared to the percentages calculated from the CD measurements on proteins in isolation. We assume that the C-termini of PfELC as well as TgELCs are partially disordered, although to a different degree, and constitute the typical calmodulin fold only upon binding to the MyoA neck domain. This seems plausible because no structure of essential light chain has been solved in isolation so far and it has been suggested that they are disordered in the unbound state [36].

The main role of essential light chains is to support the structure of the myosin neck which, together with the light chains and myosin converter domain, serves as myosin lever arm [27,36–38]. Contrary to previously published results [14], we were able to show that ELCs bind to the same conserved region of MyoA and therefore we expect the structure of PfELC bound to PfMyoA to be similar to the *T. gondii* structures presented here. We could show that the C-terminus of myosin A is disordered (TgMyoA) or only partially ordered (PfMyoA), whereas in the trimeric complexes, TgMyoA forms a continuous α helix. Therefore, we propose that the binding of essential light chains and their respective myosin binding sites mutually induces folding of both components and thereby stiffens the myosin lever arm. In turn, the myosins are capable of undergoing a larger step size and thus increase their speed, which is in agreement with the previous measurements of both *T. gondii* and *P. falciparum* myosin A motors [14,16,17].

### Implications on myosin A regulation

In general, the myosin light chains together with the myosin heavy chain neck region constitute a regulatory domain, which influences the biochemical and mechanical properties of myosins either upon phosphorylation [39–42] or by direct binding of calcium [43, 44]. Indeed, apicomplexan invasion is accompanied by an increase of intracellular calcium concentration and activation of several kinases [45–47]. The apicomplexan MLC1 or MTIP do not contain the N-terminal signature sequence (RxxS) necessary for the recognition by myosin light chain kinases as in muscle myosins [48] and therefore must follow different regulatory pathways. It has been shown that phosphorylation of TgMyoA S21 and S743 [49] and PfMyoA S19 [24] upregulates the myosin motor. On the other hand, *P. falciparum* MTIP phosphorylation at S108 impairs the affinity of MTIP to PfMyoA by increased electrostatic repulsion from the adjacent MTIP residue E179 [22]. The previously identified phosphorylation sites of MLC1 (S55, T98, S132) [47] are not homologous to MTIP phosphorylation sites (S47, S51, S85/86, S108) [22, 46] and all of them locate to the N-terminal disordered domain or distant from the binding interface with ELCs and MyoA, making them unlikely to be directly involved in myosin motor regulation. The structure of the glideosome trimeric complex 2 revealed another possible site of the myosin A regulation at TgELC2 residue S102, which has been previously shown to be phosphorylated in *T. gondii* tachyzoites [50]. In the trimeric complex, S102 forms a hydrogen bond with TgMyoA E787 and we expect that an additional negative charge would have a repulsive effect and decrease the binding affinity of TgELC2. Interestingly, residue S102 is not conserved in TgELC1 and could explain the presence of two essential light chains in *T. gondii*: they might be used by the parasite in different life stages, where a different type of myosin regulation is required. In addition, it cannot be excluded that the essential light chains are promiscuous and might be able to bind to other myosins, too. Although both *T. gondii* essential light chains have been shown to interact with the glideosome *in vivo* [16], transcriptomic data indicate a higher gene expression level of TgELC1 compared to TgELC2 throughout all life stages except during oocyst development [51], suggesting that TgELC2 might play additional roles in other cellular processes then invasion and motility.

Another mode of regulation described in classical myosins is calcium binding. Similarly, to other essential light chains, TgELC1 and TgELC2 bind calcium in their first EF hand by residues just adjacent to the interface with MLC1. The calcium ion is supposed to support the stiffness of the myosin helix kink (TgMyoA residues 801-803) and mediate transfer of the small conformational changes of the distal regulatory light chain upon phosphorylation to the myosin motor domain [35, 43]. Contrary to that, the calcium-free crystal structure of complex 1 displayed an identical conformation, forming an even larger interface with MLC1 than the calcium-bound structure. Moreover, the normal mode analysis has shown that conformational flexibility of the helical hinge of TgMyoA (residues 801-803) is limited and further propagates downstream of residues 799 with a main deformation hinge in the TgMyoA helical hinge between residues 773-777. This suggests that calcium does not play a role in myosin A regulation but rather stabilizes the complex *per se*. Such conclusion is also supported by previously published functional data, which show that the absence of calcium does not alter the function of the myosin A motor in neither *P. falciparum* [17] *n*or *T. gondii* [13].

### Assembly of the glideosome

Finally, essential light chains generally interact with the myosin converter domain and presumably stabilize the hinge of the myosin neck between the ELC and the converter domain (TgMyoA residues 775-777) [34, 35]. A small interaction interface between the converter domain and TgELC1 has also been shown previously [15]. Our models now show that both TgELC1 and TgELC2 form few polar interactions with the converter domain, however, these are not sufficient to maintain the rigid structure and the TgMyoA hinge between ELC and the converter domain contributes to most of the movement of the myosin complex. Nevertheless, the normal mode analysis was performed in the absence of a bound nucleotide and the interface between TgELCs and the converter domain might become more rigid once TgMyoA binds ATP, as previously described for other myosins [52].

It still remains elusive how myosin A and its light chains attach to the inner membrane complex and bind the glideosome associated proteins. Although MLC1/MTIP are lipidated at their N-terminus, they have been shown to form a pre-complex with GAP45 which is indispensable for correct MyoA location [10]. Given the disordered nature of the MLC1/MTIP N-termini and GAP45 in isolation [53], we propose that the proteins interact and mutually induce their secondary structure upon binding, similarly to the ELC-MyoA interaction described here. We expect that this interaction can be fine-tuned by phosphorylation of both MLC1/MTIP and GAP45. Moreover, as both GAP45 and MLC1/MTIP are lipidated and on the other hand, the transmembrane regions of both GAP40 and GAP50 are highly conserved, we propose that GAP40 or GAP50 directly bind the lipidation moieties of GAP45 and/or MLC1/MTIP to immobilize the glideosome in the 2D plane of the inner membrane complex.

In conclusion, our study extends the knowledge on the structure and mechanism of the glideosome. Partially disordered ELCs stiffen the MyoA lever arm by mutual induction of structure upon binding and together with MLC1/MTIP form complexes that are structurally conserved between *T. gondii* and *P. falciparum*. The structures revealed that calcium is not directly involved in the myosin motor regulation, but uncovered an additional potential site of regulation by phosphorylation in TgELC2. Nonetheless, the complete understanding of the molecular basis of apicomplexan motility and invasion will require further research and it will be interesting to unravel how the glideosome anchors to other proteins of the IMC and how the parasites regulate the action of differently localized myosins on a temporal scale.

## Materials and methods

### Cloning

Open reading frames encoding TgELC2 (TGME49_305050) and TgMLC1 (TGME49_257680) subcloned *via* NdeI/XhoI restriction enzymes into pET28a(+)-TEV vector were purchased from *GenScript*. The TgELC1 gene was cloned by extending the TGME49_269442 open reading frame (*GenScript*) and cloning into a pNIC28_Bsa4 [54] vector *via* BsaI restriction sites. DNA sequences of PfELC (PF3D7_1017500), PfELC-N (residues 1-74), PfMTIP (PF3D7_1246400) and PfMTIP-S (residues 60-204) were amplified from *P. falciparum* 3D7 cDNA and cloned into a pNIC28_Bsa4 vector *via* BsaI restriction sites. These constructs have an N-terminal TEV-cleavable His_6_-tag. TgMLC1-S (residues 66-146) was subcloned into a pNIC_CTHF [54] vector *via* the BfuI restriction site. The vector has a C-terminal TEV-cleavable His_6_-tag and FLAG-tag. The sequence encoding TgMyoA-C was amplified by two complementary primers and cloned *via* NcoI/KpnI restriction enzymes into a pET_GB1 vector. This construct contains an N-terminal TEV-cleavable His-GB1 domain. Expression cassettes of His-TgELC1 and His-GB1-TgMyoA-C were then subcloned *via* NdeI/XbaI restriction enzymes into a pPYC [55] vector. The His-GB1-TgMyoA-C gene was then cut by SpeI/XbaI restriction enzymes and inserted into SpeI-cut pPYC-His_TgELC1 to construct the co-expression vector pPYC with TgELC1 and TgMyoA-C.

### Mutagenesis

Site directed mutants were generated by blunt-end PCR. Briefly, the plasmids were amplified by primers which contain the alternative bases on their 5’ ends and anneal upstream and downstream of the target triplet. The PCR products were digested by DpnI (NEB) overnight at 37 °C and purified by a PCR purification kit (Qiagen). Subsequently, the 5’ ends of the PCR products were phosphorylated by T4 polynucleotide kinase (NEB), the products were purified and the free ends of the plasmid re-ligated by T4 DNA ligase (NEB). The positive clones were subsequently selected and their sequence was verified by sequencing.

### Protein expression and purification

The proteins were overexpressed in *E. coli* BL21(DE3) (MLC1, MTIP, MTIP-S, co-expressed TgELC1-TgMyoA-C + MLC1-S) or *E. coli* BL21-CodonPlus(DE3)-RIL (TgELC1, TgELC2, PfELC, PfELC-N, MLC1-S), in TB medium. The bacterial cultures were induced at OD_600nm_ of 0.6 with 1 mM IPTG and harvested after 4 hours at 37 °C (TgELC1, TgELC2, PfMTIP) or induced at OD_600nm_ of 0.6 by 0.2 mM IPTG and harvested after 16 hours at 18 °C (PfELC, PfELC-N, MLC1). The expression of PfELC and PfELC-N for NMR measurements was performed in minimal expression medium as described elsewhere [56].

The cell pellets were resuspended in lysis buffer (20 mM NaP (pH 7.5), 300 mM NaCl, 5% glycerol, 15 mM imidazole, 5 units/ml DNase I, 1 tablet of protease inhibitors (*Roche*) per 100 mL buffer, 1 mg/mL lysozyme, 0.5 mM TCEP) and the bacteria were lysed by three passages through an emulsifier (EmulsiFlex-C3, Avestin) with a maximum pressure of 10 000 psi. The lysate was centrifuged (20 min, 19 000*g*) and incubated with 2 ml of Ni-IMAC beads (ThermoFisher) per 1 l of culture on a rotatory wheel (1 h, 4 RPM). The lysate was then transferred into a gravity column and washed twice with 10 ml wash buffer (20 mM NaP (pH 7.5), 300 mM NaCl, 5% glycerol, 15 mM imidazole, 0.5 mM TCEP). The bound protein was eluted with 10 ml and subsequently with 5 ml of elution buffer (20 mM NaP (pH 7.5), 150 mM NaCl, 5% glycerol, 250 mM imidazole, 0.5 mM TCEP). The elution fractions were pooled and 0.5 mg of TEV protease per liter of bacterial culture was added. The samples were dialyzed (2 kDa cut-off) against 500 ml wash buffer or, in case of PfELC and PfELC-N, against 50 mM Tris (pH 8.0), 20 mM NaCl, 0.5 mM TCEP overnight. Next day, the samples were incubated on a gravity column with 1 ml Ni-beads per 1 l of culture. The flow-through was concentrated (10 kDa cut-off) to maximum of 10 mg/ml and further purified by size exclusion chromatography on a Superdex 200 HiLoad column (GE Healthcare; PfELC, MTIP, MTIP-S, MLC1, MLC1-S) or on a Superdex 75 HiLoad column (GE Healthcare; TgELC1, TgELC2, PfELC-N, co-expressed TgELC1-TgMyoA-C), using gel filtration buffer (20 mM HEPES (pH 7.5), 150 mM NaCl, 0.5 mM TCEP). Finally, the samples were concentrated (10 kDa cut-off) up to 15 mg/ml and either directly used or flash-frozen for later use. Due to instability, PfELC was always directly used within 3 days of the purification without freezing. All steps were performed at 4 °C.

### SDS-PAGE analysis

The concentrated samples of PfELC were dialyzed against 50 mM Tris (pH 8.0), 20 mM NaCl, and 0, 0.25, 0.5 or 1 mM TCEP overnight at 4 °C. Subsequently, the protein concentration was adjusted to 1 mg/ml and 50 μl of each sample was mixed with a fivefold excess of 2-iodoacetamide. The samples were incubated for 1 h at 37 °C and afterwards, 10 μl of each sample was mixed with 5 μl of non-reducing loading dye. The gel was run at 180 V for 40 min and stained by Direct Blue.

### Analytical gel filtration

The proteins and protein complexes were analyzed by analytical gel filtration using a Superdex 200 5/150 column (GE Healthcare) and the 1260 Infinity Bio-inert high-performance liquid chromatography system (Agilent Technologies) at 10 °C. The system and column were equilibrated in 20 mM HEPES (pH 7.5), 150 mM NaCl, 0.5mM TCEP and 30 μl of each sample was injected by an autosampler. The system was run at 0.2 ml/min for 20 minutes and the elution profile was recorded by a UV detector.

### Thermal shift assay

The stability of the different proteins was measured by nanoDSF (Prometheus NT.48, NanoTemper Technologies, GmbH). The proteins were first dialyzed against 1 l of gel filtration buffer supplemented with 5 mM EDTA overnight at 4 °C and subsequently 2x against 1 l of gel filtration buffer without EDTA overnight at 4 °C. The protein concentration was then adjusted to 100 μM (individually or 100 μM each component of a complex) in gel filtration buffer and varying concentration of calcium chloride (0 – 500 μM). 10 μl of samples was loaded in the glass capillaries and heated from 20 °C to 95 °C with a heating rate of 1 °C/min. The fluorescence signals with excitation wavelength of 280 nm and emission wavelengths of 330 and 350 nm were recorded and the melting temperature was calculated as either maximum of the derivative of ratio of fluorescence at 330 and 350 nm, or as maximum of the derivative of the fluorescence at 330 nm.

### Circular dichroism

To estimate the secondary structure content of the proteins and peptides, we measured circular dichroism on a Chirascan CD spectrometer (Applied Photophysics). For spectrum measurements, the protein or peptide concentration was adjusted to 100 μM and diluted tenfold by 10 mM NaP (pH 7.5), 20 mM NaCl, 0.25 mM TCEP just prior to the measurement. To measure the difference in secondary structure content in presence or absence of calcium, the proteins were first dialyzed against 1 l of gel filtration buffer supplemented with 5 mM EDTA overnight at 4 °C and subsequently 2x against 1 l of gel filtration buffer supplemented with ±1 mM CaCl_2_ overnight at 4 °C. The proteins were then diluted to 5 μM or 10 μM with 10 mM NaP (pH 7.5), 20 mM NaCl, 0.25 mM TCEP and ±1 mM CaCl_2_ just prior to the measurement. The CD spectrum was measured between 200 nm and 260 nm with 1 nm steps in triplicates using a 2 mm quartz cuvette. To assess the induction of structure in the dimeric protein complexes, each component was diluted by 10 mM NaP (pH 7.5), 150 mM NaF and 0.25 mM TCEP to a final concentration of 5 μM. The circular dichroism was measured 10x between 195 nm and 260 nm with 0.5 nm step in 1 mm quartz cuvette. The data were averaged, background subtracted and analyzed by K2D algorithm [57] using DichroWeb [58].

### Isothermal titration calorimetry

To measure the interaction of TgELC1 or TgELC2 and its mutants with the TgMyoA-C^ELC^ peptide (S777-Q798), the peptides were dissolved and the proteins were dialyzed in gel filtration buffer, overnight at 4 °C and 2 μl of a 200 μM peptide solution were injected 19 times into 20 μM protein. To measure the interaction of the trimeric complex, first, the peptides were dissolved and the proteins dialyzed against gel filtration buffer supplemented with 1 mM CaCl_2_. The complex of TgELC1, TgELC2 or MTIP-S with the MyoA peptide (S777-V818 in *T. gondii*, V775-V816 in *P. falciparum*) was first formed in 1:1.1 molar ratio, respectively, and incubated for 1 h at 4 °C. For measurement, 2 μl of 200 μM TgMLC-S or PfELC was injected 19 times into 20 μM of the pre-formed complex. The measurements were performed with a MicroCal PEAQ-ITC (Malvern) at 25 °C. The data were processed using the MicroCal PEAQ-ITC Analysis Software and fitted with a one-site binding model.

### Bioinformatics methods

The homologous protein sequences were aligned with the program MAFFT [59]. The protein disorder probability was calculated using the disEMBL [60] server with loops and coils defined by dictionary of secondary structure of proteins [61]. The secondary structure prediction of PfELC, TgELC1 and TgELC2 was calculated in JPred [62].

### Small angle X-ray scattering

The SAXS data were collected at the P12 BioSAXS beamline [63] at the PETRA III storage ring (DESY, Hamburg, Germany). The concentrated samples of TgELC2 and PfELC (10 mg/ml) were dialyzed against the buffer (20 mM HEPES (pH 7.5), 150 mM NaCl, 0.5 mM TCEP for TgELC2; 20 mM Tris (pH 8.0), 150 mM NaCl, 0.5 mM TCEP for PfELC-N) overnight at 4 °C. Further, the samples were centrifuged (5 min, 15 000*g*, 4 °C) and a dilution series of each sample (typically in a range of 0.5 – 10 mg/ml) and their corresponding solvent were measured at room temperature under continuous flow with a total exposure of 1 s (20 x 50 ms frames). The dimeric complex TgELC2/TgMyoA-C, as well as the trimeric complexes using different constructs, were mixed in 1:1 or 1:1:1 molar ratio, purified by SEC and concentrated to 10 mg/ml prior to the measurement. The X-ray scattering data were measured in an on-line SEC-SAXS mode, using a SD200 Increase column (GE Healthcare) at 0.5 ml/min with 1 frame recorded per second. The sample of PfELC was concentrated to 10 mg/ml and the X-ray scattering was measured in the on-line SEC-SAXS mode, using a SD200 5/150 column at 0.4 ml/min. The automatically processed data were further analyzed using the ATSAS suite [64] programs CHROMIXS [65] and PRIMUS [66] to determine the overall parameters and distance distribution, CRYSOL [67] to compute the scattering from the crystal structures and CORAL [68] to compute the scattering from the crystal structures with dummy residues mimicking the missing flexible parts.

### NMR

All NMR experiments were conducted on a Bruker Avance II 800 NMR spectrometer equipped with a cryoprobe at 288 °K in 50 mM HEPES, 20 mM NaCl, 0.5 mM TCEP and 10% (v/v) D_2_O at pH 7.0, except for H(CCO)NH-TOCSY and (H)C(CO)NH-TOCSY experiments that were performed on a Bruker Avance III 600 NMR spectrometer equipped with a room temperature probe. Full-length PfELC (residues 1-134) was ^15^N and ^15^N^13^C labeled and concentrated to 500 μM. PfELC-N was also ^15^N and ^15^N^13^C labeled and in addition site-selectively ^13^C labeled [69–71] by using 1-^13^C_1_ and 2-^13^C_1_ glucose. Samples were concentrated to about 1 mM. All spectra were processed suing NMRPipe [72] and analyzed using NMRView [73].

Backbone resonances of ^15^N^13^C labeled samples (1-74 and 1-134) were assigned using HNCACB [74] and HN(CO)CACB [75] experiments. Aliphatic side chains (1-74) were assigned using H(CCO)NH-TOCSY [76] (H)C(CO)NH-TOCSY and H(C)CH-TOCSY [77] experiments. Aromatic side chains (1-74) were assigned by (HB)CB(CGCD)HD [78] and aromatic H(C)CH-TOCSY experiments and verified by the site-selective ^13^C labeling.

NOEs for the structure determination were derived from 3D-NOESY-HSQC experiments for ^15^N, ^13^C aliphatic nuclei and ^13^C aromatic nuclei (on 1-^13^C_1_ and 2-^13^C_1_ glucose labeled samples). Phi-Psi dihedral angle constraints were derived using TALOS [79]. Structure calculations were performed using ARIA 2.3 [80] and standard parameters. The lowest-energy models have been deposited in the PDB with accession number 6tj3.

{^1^H}-^15^N NOE saturation was performed using a train of shaped 180° pulses in a symmetric fashion [81–83] for 3 s and a total inter-scan relaxation period of 10 s. Data collection, processing and analysis details are summarized in S3 Table.

### Crystallization

PfELC-N was concentrated (5kDa cut-off) to 26 mg/ml and 200 nl of the sample was mixed with 100 nl of reservoir solution (0.1M Tris-HCl (pH 8.5), 0.2M Li_2_SO_4_, 30% PEG 4000). The crystals grew in sitting drop plates at 19 °C for 7 days.

The trimeric complex of MLC1-S, TgELC2 and TgMyoA-C (S777-V818) was mixed in a molar ratio of 1.1: 1.1: 1, respectively. After 1 h of incubation, the trimeric complex was separated by gel filtration in 20 mM HEPES pH 7.5, 150 mM NaCl, 0.5 mM TCEP using a Superdex 75 16/600 column (GE Healthcare). The fractions containing the peak of the trimeric complex were concentrated (5 kDa cut-off) to 10 mg/ml. The crystals grew for 7 days at 19 °C in sitting drop plates prepared by mixing 200 nl of the sample with 100 nl of reservoir solution (0.1 M imidazole, 0.1 M MES monohydrate pH 6.5, 20% v/v PEG 500 MME, 10% w/v PEG 20 000, 0.12 M 1,6-hexadiol, 0.12 M 1-butanol, 0.12 M 1,2-propanediol, 0.12 M 2-propanol, 0.12 M 1,4-butanediol, 0.12 M 1,3-propanediol).

The recombinantly expressed dimeric complex of TgELC1 and TgMyoA-C (S777-V818) was mixed with MLC1 in 1:1.1 molar ratio, incubated for 1 h and the trimeric complex was separated by gel filtration in 20 mM HEPES pH 7.5, 150 mM NaCl, 0.5 mM TCEP using a Superdex 75 16/600 column (GE Healthcare). The fractions containing the peak of the trimeric complex were concentrated (5 kDa cut-off) to 10 mg/ml. The crystals of calcium-bound complex grew 7 days at 19 °C in a sitting drop plates prepared by mixing 200 nl of the sample with 100 nl of reservoir solution (20% w/v ethylene glycol, 10% w/v PEG 8000, 0.1M Tris (base), 0.1M bicine pH 8.5, 0.09 M sodium nitrate, 0.09 M sodium phosphate dibasic, 0.09 M ammonium sulfate). The crystals of calcium-free complex grew 7 days at 19 °C in a sitting drop plate prepared by mixing 200 nl of the sample with 100 nl of reservoir solution (32% w/v PEG 8000, 0.1M Tris pH 7.0, 0.2M LiCl).

### Data collection and structure determination

The diffraction data of the trimeric complexes were collected at the P13 EMBL beamline of the PETRA III storage ring (c/o DESY, Hamburg, Germany) at 0.966 Å wavelength and 100 °K temperature using a Pilatus 6 M detector (DECTRIS). The diffraction data of PfELC-N were collected at the P14 EMBL beamline of the PETRA III storage ring (c/o DESY, Hamburg, Germany) at 0.966 Å and 100 °K temperature using an EIGER 16 M detector (DECTRIS). The diffraction data were processed using XDS [84], merged with Aimless [85] and the phases were obtained by molecular replacement with Phaser [86], using the structure of peptide-bound TgMLC1 (PDB ID 5vt9) as a search model in case of the trimeric complexes and NMR structure as search model in case of PfELC-N. In all cases, the models were further built and refined in several cycles using PHENIX [88], Refmac [89] and *Coot* [90]. Data collection and refinement statistics are summed up in the Table 2. PyMOL was used to generate the figures, measure the angle of the helical kink, inter-molecular angles, distances and RMSDs. PDBePISA [91] was used to characterize the intermolecular interfaces. The atomic coordinates and the structure factors have been deposited in the PDB with accession numbers 6tj4, 6tj5, 6tj6 and 6tj7.

### Modelling

The modelling procedure was performed in Modeller version 9.18 [92]. We built 50 models for the TgMyoA residues 772-791. These 50 models were fused to the structure of TgMyoA (PDB ID 6due; residues 33-771). All 50 models were tilting along the bond/dihedral angle between residue 771 and the first modelled residue, that is 772; at the same time, the residues 33-771 of the 6due structure remained fixed. Thus, each of the produced models consisted of an intact crystal structure 6due (till residue 771) and *de novo* modelled fragment of 772-791. Restraints in a form of i-i+4 h-bonding pattern were imposed in order to ensure that all 50 models have an α-helical conformation along the whole length of the *de novo* modelled fragment, and also at the junction between residues 771 and 772. The crystal structure of complex 1 (PDB ID 6tj5) or complex 2 (PDB ID 6tj7) were superposed on the 50 models using the TgMyoA residues 780-791. After superposition, the modelled conformation of this fragment was removed from the merged structures, which produced models consisting of an intact crystal structure of TgMyoA (PDB 6due), the modelled helix of TgMyoA (residues 772-779) and the intact crystal structure of the complex 1 (50 models) or complex 2 (50 models), starting from the TgMyoA residue 780 of these structures. Next, all reconstructed complexes were screened against the existence of atomic clashes using the Chimera software [93] and the best five models (both complex 1 and complex 2) were energy minimized by executing 1000 steps of conjugate gradient energy minimization in the NAMD program [94]. All energy minimizations were performed in a water box with ions.

#### Normal mode analysis

Normal mode analysis (NMA) [95] was used to probe essential dynamics of the reconstructed trimeric models. The NMA was performed in an all-atom representation on the best five energy-minimized models using the BIO3D software [96]. The deformation analysis was performed, using the first 20, 50 and 100 modes, and also on the first 10 modes separately. This allowed us to not only identify possible hinge points within the studied structures of trimeric complexes, but also to determine which hinges correspond to which modes.

## Acknowledgments

We thank the Sample Preparation and Characterization facility of EMBL Hamburg for support with nanoDSF, ITC measurements and with protein crystallization. We acknowledge all group members for continuous support and feedback on the project and during manuscript preparation. We would like to thank the group of Thomas R. Schneider at EMBL Hamburg for access to the EMBL beamlines P13 and P14 and Guillaume Pompidor and Grzegorz Chojnowski for help with data processing and initial model building.

## List of abbreviations

Complex 1: trimeric complex of TgELC1, TgMyoA-C and MLC1-S with bound calcium ion
Complex 1f: trimeric complex of TgELC1, TgMyoA-C and MLC1-S, calcium-free
Complex 2: trimeric complex of TgELC2, TgMyoA-C and MLC1-S with bound calcium ion
ELC: essential light chain
GAC: glideosome associated connector
GAP: glideosome associated protein
GAPM: glideosome associated protein with multiple membrane spans
IMC: inner membrane complex
MIC1: microneme protein 1
MLC1: myosin light chain 1
MLC1-S: short construct of protein myosin light chain 1, residues 66-210
MTIP: myosin A tail interacting protein
MTIP-S: short construct of protein myosin A tail interacting protein, residues 60-204
MyoA: myosin A
SAXS: small angle X-ray scattering
SEC: size exclusion chromatography
TgMyoA-C: C-terminal construct of *T. gondii* myosin A, residues

## Supporting information captions

**S1 Figure. A** Disorder probability prediction calculated by the disEMBL server shows differences between PfELC and TgELCs, predominantly in the C-terminal region of the sequence. The disorder for the prediction was defined by dictionary of secondary structure probabilities. The amino acid residues with disorder probability above the threshold (dashed line) are predicted to be disordered. **B** Recorded SAXS curves of isolated PfELC and TgELC2 indicate conformational differences of these homologous proteins. **C** The dimensionless Kratky plot shows that TgELC2 is more compact than PfELC. Although none of the plots converges to zero, the maximum of TgELC2 is considerably closer to *sR_g_ =* √3 compared to PfELC. **D** Elution profile of on-line SEC-SAXS measurement of PfELC using a Superdex 200 5/150 column with the region used for the analysis highlighted in grey. **E** Overlay of ^15^N HSQC spectra of full-length PfELC (red) and its N-terminal construct PfELC-N (black), indicating that the construct PfELC-N is identical to the N-terminal domain of full-length PfELC. Assigned resonances are labeled. **F** Dimer of PfELC-N formed by a cysteine bond between two symmetry related molecules. **G** SDS-PAGE gel with PfELC-N samples dialyzed against buffers with varying concentration of TCEP and subsequently alkylated by 2-iodoacetamide. The results show that the protein is monomeric with the concentration of TCEP used for its biophysical characterization.

**S2 Figure.** ITC binding isotherm of PfMyoA-C titrated into MTIP measured at 25°C.

**S3 Figure. A** Small Angle X-ray scattering profiles of TgELC2 (blue) and in complex with TgMyoA-C (black) show conformational changes upon interaction. **B** Elution profile of on-line SEC-SAXS measurement of TgELC2 using a Superdex 200 10/300 column with the region used for the analysis highlighted in grey. **C** Far-UV CD spectra of both PfMyoA-C (pink) and TgMyoA-C (violet) indicate that the unbound C-terminus of MyoA is disordered (TgMyoA) or partially disordered (PfMyoA). **D** Far-UV CD data indicate that TgELC1 induces α-helical structure in TgMyoA upon binding. The individual spectra of TgELC1 (green) and TgMyoA^ELC^ (pink) do not sum up (dotted black line) to the spectrum of their dimeric complex (black continuous line), which has more pronounced features of α-helical secondary structure with lower ellipticity at 222 nm and higher ellipticity at 195 nm. The data were collected in a 1 mm cuvette at a concentration of 5 μM of each component in 10 mM NaP (pH 7.5), 150 mM NaF and 0.25 mM TCEP at 20°C.

**S4 Figure. A** Isothermal titration calorimetry of TgELC2 mutants binding to TgMyoA-C^ELC^. Individual mutations of polar residues (E10A, F79A, S101A) of TgELC2 interacting with TgMyoA-C^ELC^ do not cause major changes in the affinity of the two components, but the double mutant TgELC2E10A+H110A shows a threefold lower affinity. **B-C** Overlay of MLC1 derived from the published dimeric complex structure in grey (PDB ID 5vt9, orange) with the protein chains of the trimeric complex structures (complex 1 on the left and complex 2 on the right) shows that MLC1 does not undergo any major structural changes upon TgELC1 or TgELC2 binding. Color code of MLC1 derived from the trimeric complexes according to RMSD deviation of Cα is indicated. **D** Experimental small angle X ray scattering curves of complex 2 with the short MLC1 construct (MLC1^77-210^) and full-length MLC1 (MLC1^1-210^). The calculated scattering curve computed from the crystal structure of complex 1 fits the scattering data of complex 1 with construct MLC1^77-210^ with χ^2^ = 1.04, suggesting that MLC1 residues 77-210 form a folded and rigid entity in the complex. The experimental data of complex 1 with full length MLC1^1-213^ fit the calculated scattering data from the crystal structure of complex 1 and N-terminal MLC1 residues modelled by CORAL with χ^2^ = 1.15.

**S5 Figure. A** Comparison of far-UV circular dichroism spectra of individual ELC proteins in presence and absence of calcium ions show that calcium does not significantly alter the secondary structure of the ELCs. **B** Binding isotherms of MyoA-C titrated to the TgELC2 mutant D16A (first calcium-binding EF hand) shows that calcium does not have a major influence on the affinity of TgELC2 to the myosin A neck. **C** ITC binding isotherms of MLC1 titrated to the TgELC2/TgMyoA-C pre-complex. TgELC2 or MLC1 residues forming the binding interface were mutated individually and their affinity was measured to assess the contribution to the binding interface within the trimeric complex. The measured mutants were TgELC2 mutants R17A and E22A, and MLC1 mutants K168A, Q169A and N172A. The binding affinities were in the low nanomolar range. **D** Stability dependence of the trimeric complex upon addition of increasing concentrations of calcium illustrated by the increase in T_M_ (°C). Stability data for complex one are shown on the left and for complex two on the right. The colored points are individual measurements and “**+**“ represents the average. The experiment shows that the stability of the trimeric complex is greatly enhanced by the addition of calcium in a concentration-dependent manner.

**S6 Figure.** A Summary plot of the atomic displacements predicted by NMA based on the five lowest-energy models for complex 2 selected by the lowest clash score. The relative atomic displacement of the individual amino acid residues follows the same pattern in all five models, confirming that the results of the normal mode analysis are independent of the chosen starting conformation or the energy-minimized model. The results for complex 1 are similar (data not shown). **B** The deformation analysis of complex 1 averaged through the 20 lowest-energy modes predicts two main hinge regions, with the hinge 1 (residues 773-777), having the largest contribution to the observed motions. The model is coloured by the deformation energy from low (violet) to high (red). **C** The ensemble of the structures of complex 2 based on the two lowest-energy modes, which contribute most to the large-scale dynamics of proteins. The original model is drawn in cartoon representation with shown semitransparent surface, whereas the deformed structures are partially transparent and drawn in ribbon representation with faded surface. The structures were aligned on MLC1 to reflect the immobilization of MLC1 in the IMC membrane as in the current model of the glideosome (see Fig 1A).

**S1 Table. A list of published *P. falciparum* and *T. gondii* glideosome protein structures published.** So far, only the structures of individual proteins of glideosome and two homologous subcomplexes (MTIP/PfMyoA and MLC1/TgMyoA) have been solved.

**S2 Table. SAXS sample details, data acquisition parameters, structural parameters and atomistic modelling.**

**S3 Table. Statistics for NMR structure calculation of PfELC (residues 1-74).**

Ramachandran analysis was performed by PROCHECK [97].

**S4 Table. Polar interactions in the structures of the trimeric complexes inspected by Molprobity [98].**

